# The lethal triad: SARS-CoV-2 Spike, ACE2 and TMPRSS2. Mutations in host and pathogen may affect the course of pandemic

**DOI:** 10.1101/2021.01.12.426365

**Authors:** Matteo Calcagnile, Patricia Forgez, Marco Alifano, Pietro Alifano

## Abstract

Variants of SARS-CoV-2 have been identified rapidly after the beginning of pandemic. One of them, involving the spike protein and called D614G, represents a substantial percentage of currently isolated strains. While research on this variant was ongoing worldwide, on December 20^th^ 2020 the European Centre for Disease Prevention and Control reported a Threat Assessment Brief describing the emergence of a new variant of SARS-CoV-2, named B.1.1.7, harboring multiple mutations mostly affecting the Spike protein. This viral variant has been recently associated with a rapid increase in COVID-19 cases in South East England, with alarming implications for future virus transmission rates. Specifically, of the nine amino acid replacements that characterize the Spike in the emerging variant, four are found in the region between the Fusion Peptide and the RBD domain (namely the already known D614G, together with A570D, P681H, T716I), and one, N501Y, is found in the Spike Receptor Binding Domain – Receptor Binding Motif (RBD-RBM). In this study, by using *in silico* biology, we provide evidence that these amino acid replacements have dramatic effects on the interactions between SARS-CoV-2 Spike and the host ACE2 receptor or TMPRSS2, the protease that induces the fusogenic activity of Spike. Mostly, we show that these effects are strongly dependent on ACE2 and TMPRSS2 polymorphism, suggesting that dynamics of pandemics are strongly influenced not only by virus variation but also by host genetic background.

## Introduction

Viruses, like all other species, obey evolutionary and biodiversity rules. According to these rules, surviving viruses adapt to their own benefit. To prevent these adaptations, there are two human interventions possible, either eradicate the virus, or attempt to understand the relationship between the host and the virus, to mitigate the effects of the virus.

The severe acute respiratory syndrome corona virus −2 (SARS-CoV-2) spread incredibly quickly between people, due to its newness and transmission route, but the kinetics of diffusion and mortality remains variable from one country to another. A multitude of factors may concur to explain the ethnic and geographical differences in pandemic progression and severity. Considerable individual differences in susceptibility to onset of disease caused by SARS-CoV-2 may be involved, including mainly sex, age and underlying conditions [1]. However, genomic predisposition, a major concept in modern medicine, prompts to understand molecular bases of heterogeneity in diffusion and severity of disease, to find prevention and treatment strategies. If host genomic predisposition has been advocated since the beginning of pandemic, virus’ genomic predisposition (i.e. the occurrence of mutations) to spread less or more easily and eventually to cause more or less severe disease has been initially unkempt, because of capacity of Coronaviruses to proofread, thus removing mismatched nucleotides during genome replication and transcription. However, in the last few months it begun evident that some specific mutants (see below) are progressively superseding the virus that was identified as wild type, rising enormous questions in terms of comprehension of mechanisms of pathogen-host interaction.

SARS-CoV-2 uses envelope Spike projections as a key to enter human airway cells [2] through a specific receptor. Spike glycoproteins form homotrimers protruding from the viral surface, and comprise two major functional domains: an N-terminal domain (S1) for binding to the host cell receptor, and a C-terminal domain (S2) that is responsible for fusion of the viral and cellular membranes [3]. Following the interaction with the host receptor, internalization of viral particles is accomplished thanks to the activation of fusogenic activity of Spike, as a consequence of major conformational changes that are triggered by receptor binding, low pH exposure and proteolytic activation [3]. Spike glycoproteins are cleaved by furin at the boundary between S1 and S2 domains, and the resulting S1 and S2 subunits remain non-covalently bound in the prefusion conformation with important consequences on fusogenicity [3]. Notably, at variance with SARS-CoV and other SARS-like CoV Spike glycoproteins, SARS-CoV-2 Spike glycoprotein contain a furin cleavage site at the S1/S2 boundary, which is cleaved during viral biogenesis [4], and may affect the major entry route of viruses into the host cell [3].

Proteases from the respiratory tract such as those belonging to the transmembrane protease/serine subfamily (TMPRSS), TMPRSS2 or HAT (TMPRSS11d) are able to induce SARS-CoV Spike glycoprotein fusogenic activity [5-8]. The first cleavage at the S1-S2 boundary (R667) facilitates the second cleavage at position R797 releasing the fusogenic S2’ sub-domain^5^. On the other hand, there is also evidence that cleavage of the ACE2 C-terminal segment by TMPRSS2 can enhance Spike glycoprotein-driven viral entry [9]. Notably, it has been demonstrated that SARS-CoV-2 cell entry is blocked by specific protease inhibitors [10].

SARS-CoV-2 and respiratory syndrome corona virus (SARS-CoV) Spike proteins share very high phylogenetic similarities (99%), and both viruses exploit the same human cell receptor namely angiotensin-converting enzyme 2 (ACE2), a transmembrane enzyme whose expression dominates on lung alveolar epithelial cells [4,11,12]. This receptor is an 805-amino acid long captopril-insensitive carboxypeptidase with a 17-amino acids N-terminal signal peptide and a C-terminal membrane anchor. It catalyzes the cleavage of angiotensin I into angiotensin 1-9, and of angiotensin II into the vasodilator angiotensin 1-7, thus playing a key role in systemic blood pressure homeostasis, counterbalancing the vasoconstrictive action of angiotensin II, which is generated by cleavage of angiotensin I catalyzed by ACE^17^Although ACE2 mRNA is expressed ubiquitously, ACE2 protein expression dominates on lung alveolar epithelial cells, enterocytes, arterial and venous endothelial cells, and arterial smooth muscle cells [13].

There is evidence that ACE2 may serve as a chaperone for membrane trafficking of an amino acid transporter B0AT1 (also known as SLC6A19), which mediates the uptake of neutral amino acids into intestinal cells in a sodium dependent manner [14]. Recently, 2.9 Å resolution cryo-EM structure of full-length human ACE2 in complex with B0AT1 was presented, and structural modelling suggests that the ACE2-B0AT1 can bind two Spike glycoproteins simultaneously [15,16]. It has been hypothesized that the presence of B0AT1 may block the access of TMPRSS2 to the cutting site on ACE2 [15,16]. B0AT1 (also known as SLC6A19) is expressed with high variability in normal human lung tissues, as shown by analysis of data available in Oncomine from the work by Weiss et al [17].

Notably, a wide range of genetic polymorphic variation characterizes the ACE2 gene, which maps on the X chromosome, and some polymorphisms have been significantly associated with the occurrence of arterial hypertension, diabetes mellitus, cerebral stroke, septal wall thickness, ventricular hypertrophy, and coronary artery disease [18-20]. The association between ACE2 polymorphisms and blood pressure responses to the cold pressor test led to the hypothesis that the different polymorphism distribution worldwide may be the consequence of genetic adaptation to different climatic conditions[20,21]. I*n silico* tools identified ACE2 single-nucleotide polymorphisms (SNPs) responsible for increased or decreased ACE2/Spike affinity, suggesting that ACE2 polymorphism can contribute to ethnic and geographical differences in SARS COVID-19 spreading across the world. While these results need biological confirmation, it has become urgent that a more precise assessment of the interplay between SARS CoV-2 Spike ACE2 and TMPRSS should be evaluated taking into account polymorphisms of these two proteins along with the genetic evolution of the virus. In the context of the present pandemic, *In silico* studies provide a rapid means to evaluate the interaction between molecules and gives direction for further biological and clinical studies.

Two Spike mutations lead the current scientific debate: D614G and N501Y, the former identified as soon as January 2020, and currently accounting for the majority of isolated strains in several countries, the latter isolated form clinical samples in England and Wales in the last few days and responsible for un “uncontrolled” diffusion on the virus. Although these mutations affect different region of the Spike (directly the Receptor Binding Domain for N501Y, and the region close to the TMPRSS proteolytic site the D614G), both are likely to affect structure and function of the protein, and, as a consequence the effectiveness of the whole process of entry of the virus, though in a different manner. To elucidate initial steps of host-pathogen interactions taking into account both Spike and human polymorphisms (of both ACE2 and TMPRSS), we studied geographical distribution of the D614G variant and its evolution over time, and modeled, by *in silico* tools, on one hand D614G Spike /TMPRSS2 interaction, and, on the other one, N501Y Spike/ACE2 interaction.

## Results

### The emerging variant of SARS-CoV-2 Spike with multiple amino acid changes, and geographical distribution of the single amino acid changes as inferred from databases

On December 20^th^ 2020 the European Centre for Disease Prevention and Control reported a Threat Assessment Brief describing the emergence of a new variant of SARS-CoV-2, named B.1.1.7, harboring multiple mutations mostly affecting the Spike protein (Fig. 1A) (European Centre for Disease Prevention and Control, 2020) [22]. This variant has been recently associated with a rapid increase in COVID-19 cases in South East England, and its spread worries all governments around the world. Seven mutations are found in this variant: the mutation N501Y is found in the Spike Receptor Binding Domain – Receptor Binding Motif (RBD-RBM) and it was predicted to increase substantially the affinity of Spike for ACE2 [23], four, A570D, D614G, P681H, T716I, are found in the region between the Fusion Peptide and the RBD domain, two, S982A and D1118Y, are located, respectively, in the Heptad Repeat region 1 (HR1) and HR1-Heptad Repeat region 2 (HR2), while two small deletions, i.e. deletion 69-70 and deletion 144, affect the N-terminal region of the Spike protein (Fig. 1AB). Here we focused on the possible effects of some of these amino acid replacements on the interactions between Spike and ACE2 or TMPRSS2, and examined the possible effects of ACE2 and TMPRSS2 polymorphisms on these interactions.

**Fig 1.**
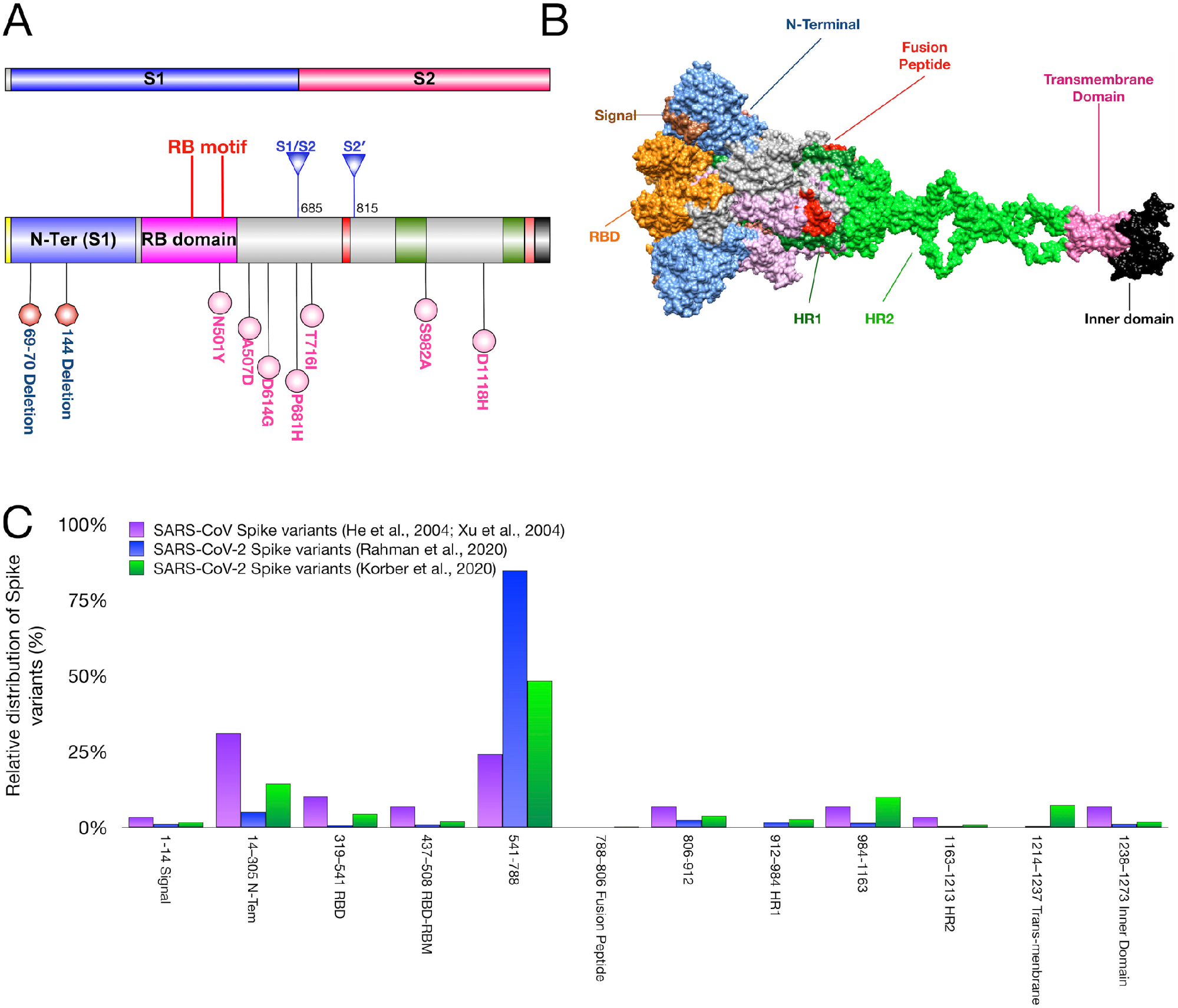
SARS-CoV and SARS-CoV-2 Spike protein variants and protein domains. **A-B)** 2D (**A**) and 3D (**B**) maps of SARS-CoV-2 Spike protein domains. Proteolytic cleavage sites (S1/S2, S2’) and amino acid variations identified in the B.1.1.7 SARS-CoV-2 variant are shown in the 2D map. **C**) Relative distribution of SARS-CoV and SARS-CoV-2 Spike amino acid variations with respect to each protein domain as inferred by Dataset S1 (SARS-CoV-2), Dataset S2 (SARS-CoV-2) and Dataset S3 (SARS-CoV). RBD, Receptor Binding Domain; RBM, Receptor Binding Motif; HR1, Heptad Repeat region 1; HR2, Heptad Repeat region 2.

On June 2020 the frequencies of the individual amino acid substitutions (N501Y, A570D, D614G, P681H, T716I, S982A, D1118Y) in COVD-19 clinical samples were quite low except the frequency of the D614G that was already high suggesting that the punctual mutation D614G represented already a selective advantage for the virus [24,25] (Table 1). Figure 1C illustrates the relative distribution of annotated SARS-CoV-2 Spike mutations with respect to each protein domain as inferred from [24] (Dataset S1), and [25] (Dataset S2), and also shows the results of this analysis with SARS-CoV Spike mutations [26,27] (Dataset S3). Spike protein variants encompassing 7 domains (N-terminal, receptor-binding domain (RBD), fusion peptide, HR1, HR2, trans-membrane domain and inner domain), and 3 inter-domain regions (RBD-fusion peptide, fusion peptide-HR1 and HR1-HR2) were analyzed. The data show that most common variants of the SARS-CoV-2 Spike are located in a protein region that spans the amino acids 541 and 788, while in SARS-CoV Spike the corresponding region between amino acids 14 and 305 was primarily affected by amino acid variation, followed by the region 541-788 (Fig. 1C). The region 541-788 links the RBD and the fusion peptide and contains the S1/S2 cleavage site for TMPRSS2, the trans-membrane protease that has been shown to carry out the priming of the SARS-CoV-2 Spike by sequential cleavage at S1/S2 and S2’ (Fig. 1C) [10,28].

**Table 1.** Frequencies of SARS-CoV-2 Spike amino acid substitutions detected in COVD-19 clinical samples.

| Amino acid substitution | Position | Domain | Number of detected allele in the database<sup>a</sup> | Relative frequency of allele (%) |
| --- | --- | --- | --- | --- |
| N501Y | 501 | RBD-RBM | 1 | 0.00% |
| A570V | 570 | Fusion Peptide-RBD | 13 | 0.04% |
| A570S | 570 | Fusion Peptide-RBD | 3 | 0.01% |
| A570T | 570 | Fusion Peptide-RBD | 1 | 0.00% |
| A570D | 570 | Fusion Peptide-RBD | 1 | 0.00% |
| D614G | 614 | Fusion Peptide-RBD | 26408 | 82.24% |
| D614N | 614 | Fusion Peptide-RBD | 4 | 0.01% |
| P681L | 681 | Fusion Peptide-RBD | 20 | 0.06% |
| P681S | 681 | Fusion Peptide-RBD | 4 | 0.01% |
| P681H | 681 | Fusion Peptide-RBD | 3 | 0.01% |
| T716I | 716 | Fusion Peptide-RBD | 14 | 0.04% |
| S982A | 982 | HR1 | 0 | 0.00% |
| D1118Y | 1118 | HR2-HR1 | 2 | 0.01% |

The database was constructed from the data in Rhaman et al., 2020 [24].

The distribution and temporal spread of the SARS-CoV-2 Spike D614G, the most diffused variant. The data show distinct patterns in the different geographical regions. Specifically, in Eastern Asia the D614G variant was described in January 2020 at low frequency (3.8%) and then it was subject to a decremental trend over time (Fig. S1). A similar trend can be also observed in Central Asia, where the variant was reported in March 2020. In contrast, an incremental trend can be noted in South Eastern Asia with a maximal occurrence approaching 30% in June 2020. The D614G variant reached the highest occurrences in Northern Western Europe (50% in April 2020), Central Europe (18% in February 2020), Southern Europe (10% in February 2020), and Northern America (26% in March 2020) with distinct temporal patterns but a persistence trend in the population. Much lower frequencies can be observed in Eastern Europe and Russia, Southern America and in Africa, with an incremental trend in Eastern Europe and Africa. The geographical distribution, the different occurrence and temporal spread of the SARS-CoV-2 Spike protein D614G variant worldwide raises the question of whether genetic differences of the host may be involved.

### Effects of D614G on SARS-CoV-2 Spike protein structure as inferred by *in silico* modeling

*In silico* simulations were performed to investigate the possible effects of the D614G substitution on SARS-CoV-2 Spike protein folding and flexibility. Secondary structures of the region spanning the amino acids 601-627 in the wild type and the D614G variant of SARS-CoV-2 Spike protein were predicted by using the *ab initio* method on PEPfold server [29,30]. The results demonstrated that in the wild type (D614) SARS-CoV-2 Spike this region forms an N-terminal α-helix followed by a C-terminal β-sheet (Fig. 2A). D614G replacement was predicted to drastically change the peptide secondary structure by replacing the C-terminal β-sheet with a α-helix (Fig. 2A). The effects of the D614G substitution on SARS-CoV Spike structure were then analyzed. Unlike the SARS-CoV-2 Spike, the corresponding amino acid region (amino acids 587-613) in the wild type SARS-CoV Spike is arranged in two anti-parallel α-helices (Fig. 2B), similar to those found in D614G SARS-CoV-2 Spike variant, and the D614G substitution does not seem to change this structure (Fig. 2B).

**Fig 2.**
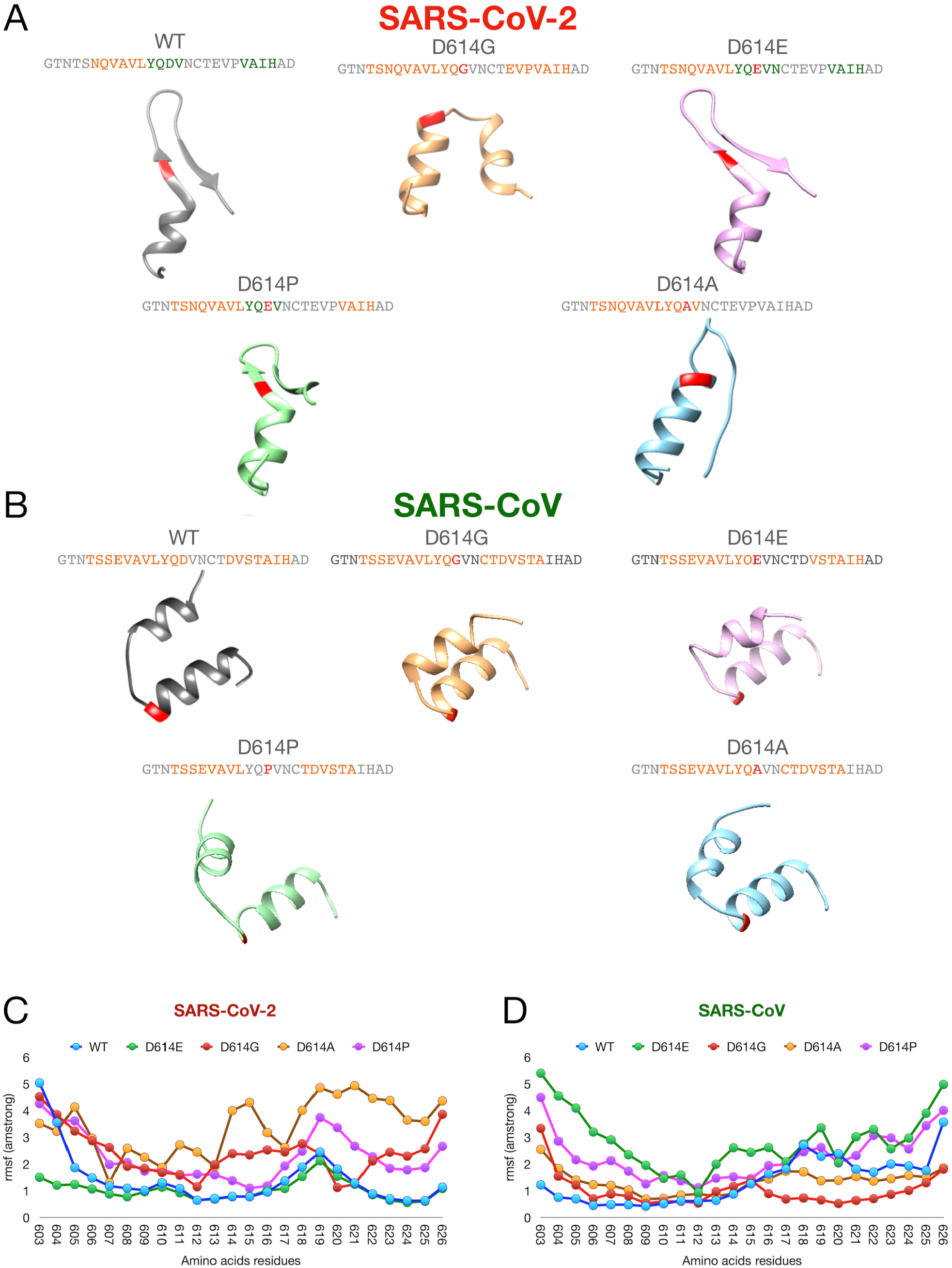
Regional secondary structures prediction and domain flexibility of wild type and variants of SARS-CoV and SARS-CoV-2 Spike proteins. **(A-B)** Secondary structures of the region spanning the amino acids 601-627 of Spike, wild type containing diverse amino acid substitutions in position 614: (**A**) SARS-CoV-2 Spike; (**B**) SARS-CoV Spike. **C)** Flexibilities of the regions spanning the amino acids 601-627 of wild type and variants of SARS-CoV Spike (left), and the corresponding region of wild type and variants of SARS-CoV Spike (right) were predicted by CABSflex simulations. RMSF, Route Means Square Fluctuation.

The analysis was then extended to other D614 variants. Results show that in SARS-CoV Spike the anti-parallel α-helices are very stable, and do not appear to be affected by any substitution studied (D614E, D614P, D614A) (Fig. 2B). In contrast, in SARS-CoV-2 Spike D614E and D614P substitution does not change the N-terminal β-sheet / C-terminal α-helix structure of the wild type protein, while in the D614A variant the C-terminal β-sheet is lost (Fig. 2A).

CABSflex [31] was used to analyze the possible effects of the D614G substitution on flexibility of the region spanning the amino acids 601-627 in SARS-CoV-2 and the corresponding region (amino acids 587-613) in SARS-CoV Spike. CABSflex simulations predicted contrasting effects in SARS-CoV-2 and SARS-CoV Spike proteins, with an increase in flexibility in the former (Fig. 2C), and a decrease in flexibility in the other protein (Fig. 2D). CABSflex analysis was also extended to the other D614 variants, and demonstrated that the increase in flexibility was highest in D614A variant (Fig. 2C), while confirming the high stability of the corresponding region (amino acids 587-613) in all SARS-CoV Spike variants (Fig. 2D).

Using PEPfold and CABSflex, a comprehensive analysis of all amino acid variations identified in the emerging B.1.1.7 SARS-CoV-2 variant was performed (Fig. S2). The computational analysis demonstrated that two amino acid substitutions, T716I and D1118H, (Fig. S2F and S2H) and the deletion at the residues 144 (Fig. S2B) may locally affect the secondary structures of the Spike protein causing the transition from β-sheet to coiled-coil structure. In contrast, N501Y, A570D and S982A substitutions (Fig. S2C, S2D and S2G) and the deletion 69-70 (Fig. S2A) did not appear to disturb the structures, while P681H (Fig. S2E) was expected to cause a minor change in the secondary structure conformation. CABSflex analysis predicted that the variations analyzed in this study generally reduce the flexibility of the Spike (Fig. S2I), particularly those affecting the amino acids 460-490 and 1050-1255. A punctual analysis revealed that the changes in amino acids 501, 716, 982, 1118 may contribute to reduce the flexibility of Spike in the B.1.1.7 SARS-CoV-2 variant, while changes in amino acids 570, 614 and 681 may have an opposite effect (Fig. S2I).

### *In silico* interaction between TMPRSS2 and wild type or D614G SARS-CoV-2 Spike, and variation with TMPRSS2 polymorphisms

The S1/S2 TMPRSS2 cleavage site was mapped at location 685 in the amino acid sequence of SARS-CoV-2 Spike protein, close to the aspartic acid residue (D614). This premise led us to explore the possible effect of the D614G substitution on TMPRSS2 processing of SARS-CoV-2 Spike protein, and a possible correlation between the geographical distribution / temporal spread of the SARS-CoV-2 Spike protein D614G variant, and the geographical distribution of TMPRSS2 polymorphisms in humans. Beside, an analysis of occurrence of residues revealed that the D614G polymorphisms was registered in both the dataset of SARS-CoV-2 (Dataset S1 and S2), in the of SARS-CoV (Dataset S3), and in a set of sequences of SARS-like viruses (Fig. S3). Bat SARS-like viruses show acid residues (D or E), while porcine and bovine SARS-like viruses show a basic residue (N), evidencing as the acid-apolar (D-G) substitution was an adaptation at the human host.

The complete list of TMPRSS2 variants was downloaded from gnomAD v2.1.1 database (https://gnomad.broadinstitute.org/), and the data were analyzed by using the multivariate ordination method of PAST. NM-MDS was used to visualize TMPRSS2 variant distribution in 7 geographical regions (African/African American, Latino/Admixed American, European (Finnish), European (non-Finnish), Ashkenazi Jewish, Southern Asian and Eastern Asian) (Fig. S4A), while PCA was used to represent the most diffused variants (>0.05% in at least one region) (Fig. S4B, Table 2).

**Table 2.**
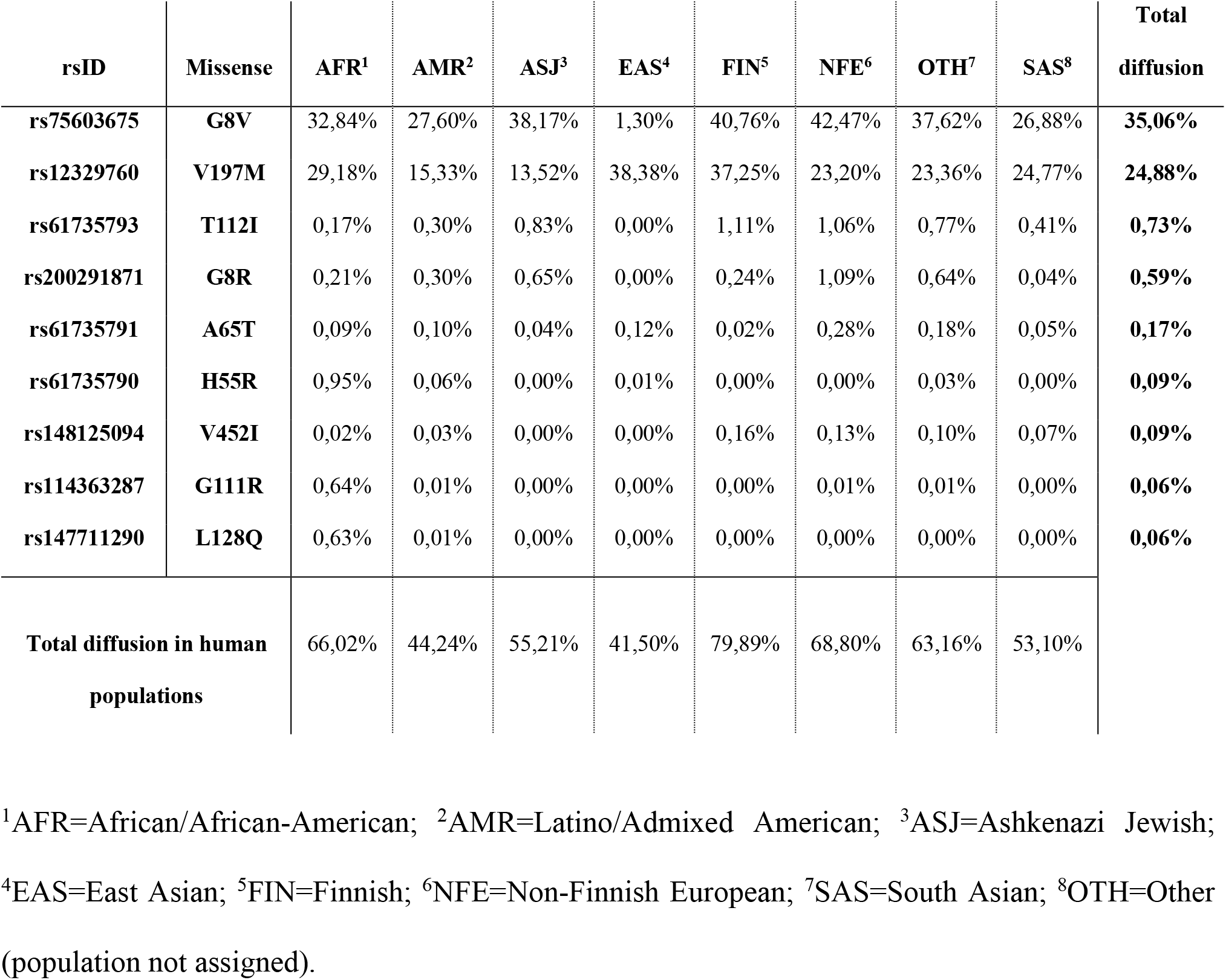
Frequencies of TMPRSS2 polymorphisms according GnomAD (frequency>0,05% for almost one measure).

Protein variants affect different regions of TMPRSS2 that contain three major functional domains: an N-terminal domain (amino acids 1-184) comprising a Low-Density Lipoprotein Receptor Class A domain (cysteine-rich repeat, LDLa, cd00112) (amino acids 150-184), a Scavenger receptor cysteine-rich domain (SRCR_2, pfam15494) (amino acids 190-283), and a Trypsin-like serine protease domain (Tryp_SPc, cd00190) (amino acids 293-524). Particularly, the N-terminal and the SRCR_2 domains exhibit the higher values of both percentages of missenses in GenomAD (n° of single missense/ total missenses) and frequencies of variants (Fig. 3A). The most diffused polymorphisms are G8V and V197M (Fig. 3B; Fig. S4C). V197M is diffused in all geographical regions (frequency >10%), while G8V is highly diffused in all regions (frequency >10%), excluding Eastern Asia (frequency <10%).

**Fig 3.**
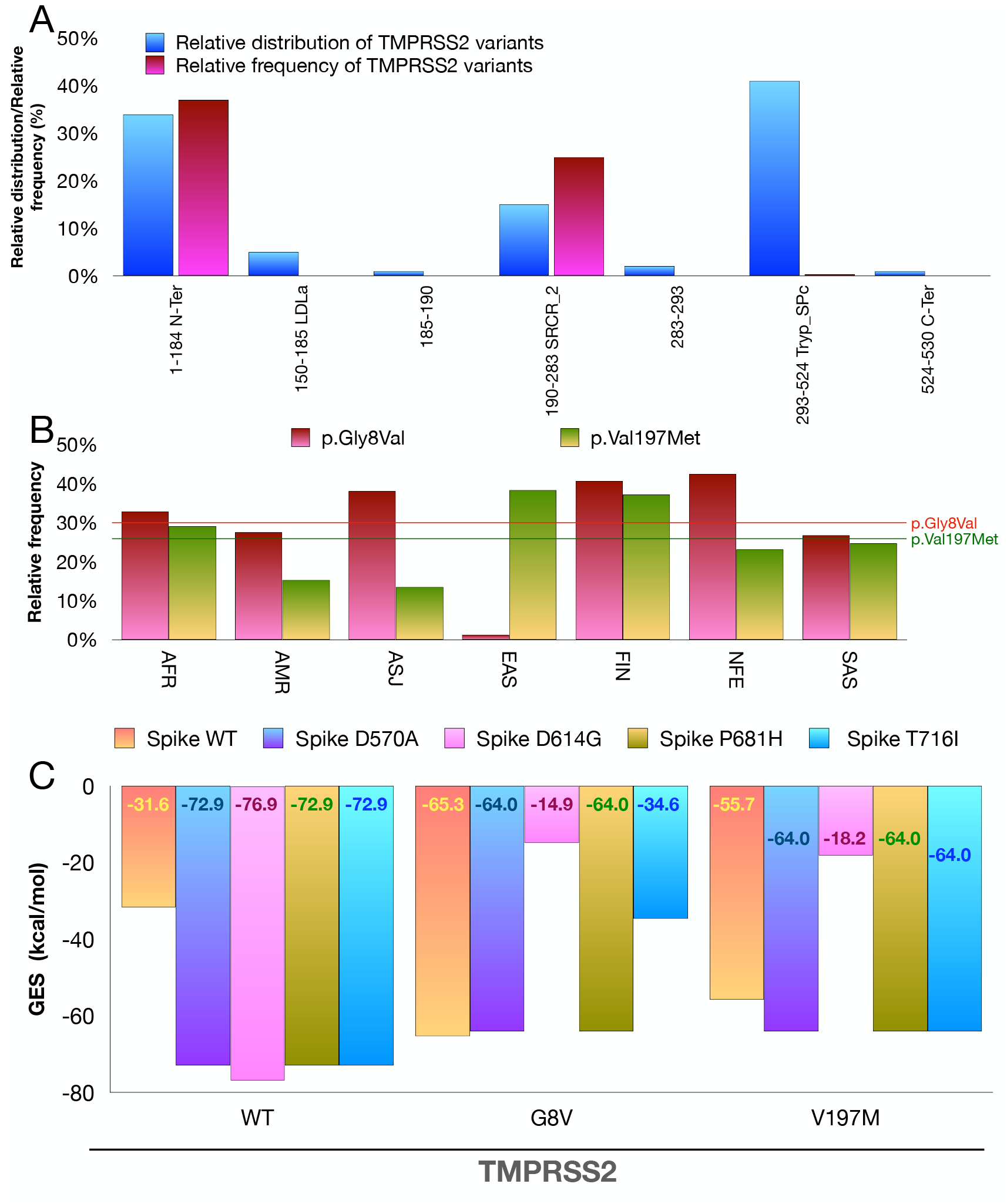
TMPRSS2 protein variants and molecular docking simulations of SARS-CoV-2 Spike protein/TMPRSS2 complexes. **A)** Relative distribution of TMPRSS2 amino acid variations with respect to each protein domain, and relative frequency worldwide according to GnomAD database. **B)** Relative frequency of G8V and V197M TMPRSS2 variants in 7 human populations. AFR, African/African-American; AMR, Latino/Admixed American; ASJ, Ashkenazi Jewish; EAS, East Asian; FIN, Finnish; NFE, Non-Finnish European; SAS, South Asian. **C)** Molecular docking simulations of SARS-CoV-2 Spike /TMPRSS2 complexes. Wild type A570D, D614G, P681H, or T716I SARS-CoV-2 Spike was used as a receptor, and wild type, G8V or V197M TMPRSS2 as a ligand. Docking simulations were carried out by Gramm-X server, and FireDock was used to calculate the GES (Global Energy Scores) as detailed in the Materials and Methods.

*In silico* molecular docking simulations were carried out on Gramm-X server [32] to predict the possible the impact of G8V and V197M TMPRSS2 variants on the interaction with either wild type or D570A, D614G, P681H and T716I SARS-CoV-2 Spike protein variants. The results demonstrated that all (D570A, D614G, P681H, T716I) substitutions result in a notable increase in the computed affinity of SARS-CoV-2 Spike protein for wild type TMPRSS2 (Fig. 3C), while a dramatic decrease in the affinity for G8V TMPRSS2 variant was observed when D614G and T716I Spike protein variants were used in the simulations (Fig. 3C). In addition, both V197M and G8V TMPRSS2 protein variants exhibited a very low affinity for D614G SARS-CoV-2 Spike.

The docking complexes obtained with wild type TMPRSS2 or G8V variant and wild type SARS-CoV-2 Spike or D614G variant were visualized by using Chimera (Fig. 4). Chimera enlightened a serine residue at position 637 of wild type SARS-CoV-2 Spike establishing an H-bond with a glutamic acid residue at position 60 of wild type TMPRSS2 (Fig. 4A). The H-bond was conserved in the complexes formed between the wild type Spike and G8V TMPRSS2 (Fig. 4B), and between D614G Spike and G8V TMPRSS2 (Fig. 4D), while it was absent in the complex formed between D614G Spike and wild type TMPRSS2 (Fig. 4C). Two additional H-bonds were present in the complexes formed between either wild type or D614G Spike and G8V TMPRSS2 (Fig. 4B and Fig. 4D, respectively): the first one involving a lysine residue at position 529 of the Spike and a glutamic acid residue at position 22 of TMPRSS2; the second one involving a phenylalanine residue at position 2 of the Spike and a glutamic acid residue at position 53 of TMPRSS2. These two H-bond were absent in the complex of D614G Spike and wild type TMPRSS2 (Fig. 4C), which showed a unique arrangement involving four H-bonds: i.) between a glutamine residue at position 271 of the Spike protein and a glutamine residue at position 66 of TMPRSS2; ii.) between a threonine residue at position 236 of the Spike protein and a threonine residue at position 68 of TMPRSS2; iii.) between a leucine residue at position 7 of the Spike protein and a serine residue at position 298 of TMPRSS2; iv.) between a threonine residue at position 20 of the Spike protein and a serine residue at position 355 of TMPRSS2. Notably, serine 298 is close to histidine 296 of the TMRSS2 active site pocket catalytic triad that also includes aspartic acid 345 and serine 441 [33].

**Fig 4.**
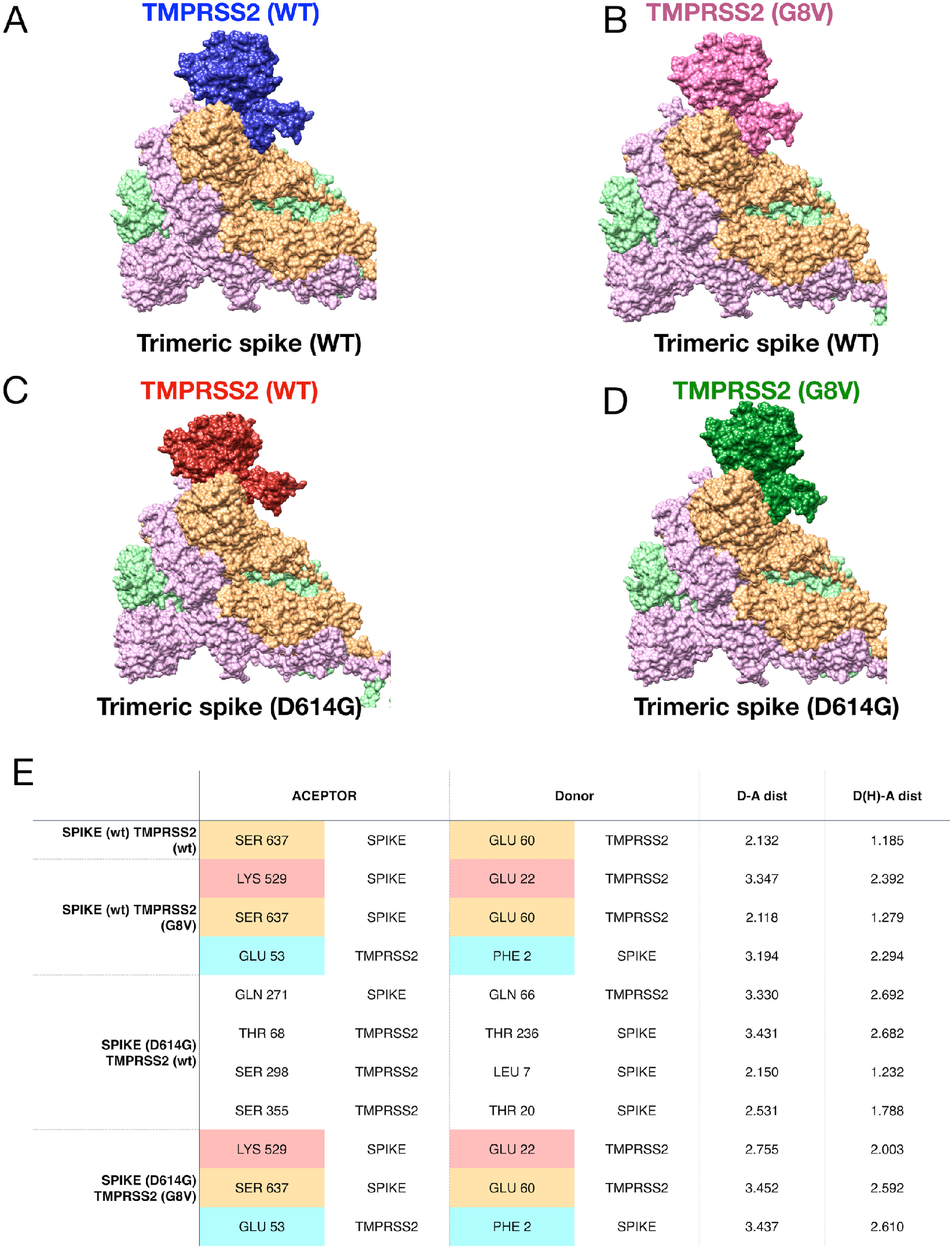
Analysis of SARS-CoV-2 Spike protein/TMPRSS2 complexes. **A-D)** Molecular docking complexes between wild type or G8V variant of TMPRSS2, and trimeric wild type or D614G variant of SARS-CoV-2 Spike were visualized by using Chimera. **E)** H-bonds analysis results. Amino acid residues providing either donor or acceptor and distances are shown (Å). D-A dist, distance between donor residue and acceptor residue; D(H)-A dist, distance between H of donor and acceptor residues.

### D614G substitution in the context of the new SARS-CoV-2 Spike variant B.1.1.7

The amino acid substitution N501Y is found in the Spike Receptor Binding Domain – Receptor Binding Motif (RBD-RBM), and this led us, in a previous work [23], to speculate possible effects on binding with ACE2. Consistently with previous finding [23], molecular docking simulations by HDOCK server predicted a substantial increase in computed binding affinity (global energy score) from −50.26 Kcal/mol (wild type Spike) to −67.49 Kcal/mol (N501Y Spike). In contrast, N501Y was expected to slightly decrease the affinity for K26R ACE2 variant (from −54.79 Kcal/mol to −52.57) (Fig. 5A), which is associated with an increased affinity with WT spike [23], suggesting that the effect of N501Y substitution is dependent on ACE2 genetic background. In fact, the analysis of molecular docking complexes by Chimera revealed that the number and the arrangement of contacts and H-bonds were different in the different complexes (Table 3), with a higher contact density in N501Y Spike RBD / wild type ACE2 complex (Fig. 5B), as compared to the other complexes. Most of contacts involve the tyrosine 501, as visualized by Chimera (Fig. 5E). Many contacts were lost in the complex between N501Y Spike RBD and K26R ACE2 variant (Fig. 5D).

**Table 3.**
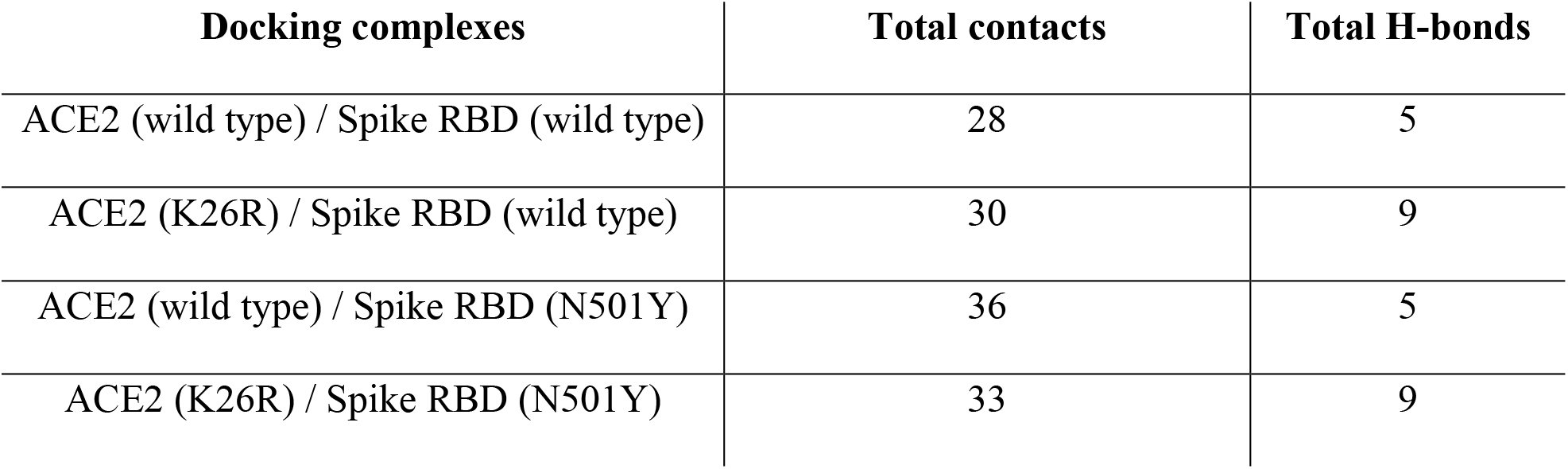
Total contacts and H-bonds in molecular docking complexes.

**Fig 5.**
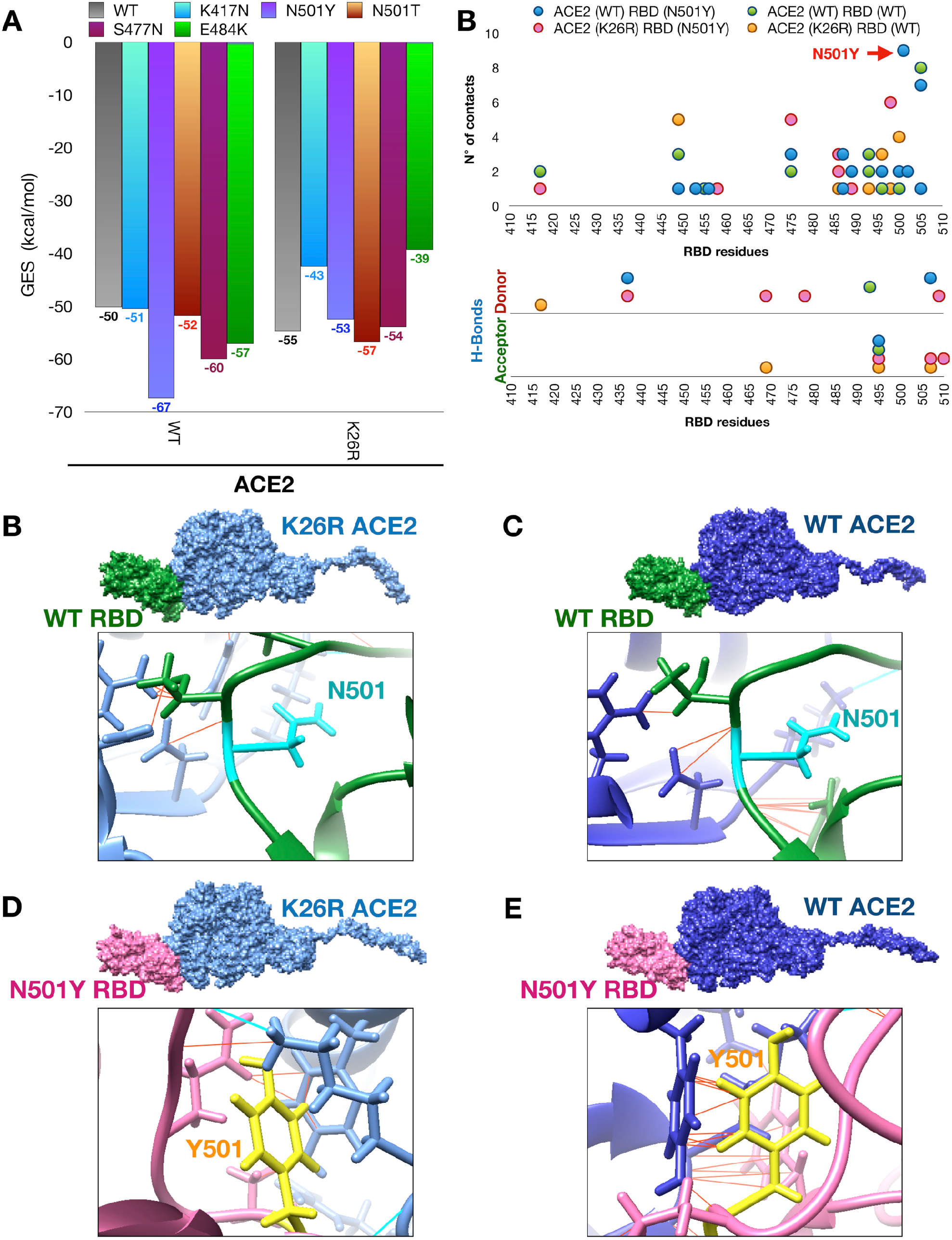
Docking complexes between wild type or K26R ACE2 and wild type or mutated SARS-CoV-2 Spike RBD. **A)** Computed affinity of molecular docking complexes between wild type wild type or K26R variant of ACE2, and wild type or mutated (N501Y, N501T, K417N, S477N, E484K) of Spike RBD. **B)** Graphical representation of the identified interaction in SARS-CoV-2 Spike RBD. Up: number of contacts (y-axis) and distribution along the RBD (x-axis). Bottom: position identified H-bonds. **C-F)** Structural analysis of docking complexes as indicated with a focus on the amino acid residue in position 501.

Figure 5A also predicts the effects of single Spike RBD-RBM amino acid substitutions, which were found in other new SARS-CoV-2 Spike variants, namely the 501.V2 variant from South Africa (K417N and E484K in addition to N501Y), the 20A.EU2 from Europe (S477N), and a new variant from Italy (harboring N501T). Results demonstrated that all RBD-RBM amino acid substitutions result in an increase in computed affinity with ACE2, albeit with substantial differences. In particular, the N501T exhibited slight increase as compared to wild type Spike (from −50.26 to −51.88 Kcal/mol), and the increase was only moderate with S477N (from −50.26 to −60.01 Kcal/mol) and E484K (from −50.26 to −57.17 Kcal/mol). Slight effects were observed with N501T and S477N using the K26R ACE2 variant as a receptor. In contrast, E484K showed a marked reduction in computed affinity with the K26R ACE2 variant (from −54.79 Kcal/mol to −39.37 Kcal/mol).

As the 501.V2 variant from South Africa includes multiple RBD-RBM amino acid substitutions affecting the Spike RBD-RBM, HDOCK simulations were then performed in the presence of all substitutions (Fig. 6A). Results predicted that the combined effect of the three amino acid substitutions N501Y, K417N and E484K was less than that of the single N501Y (Fig. 5A) in terms of increased computed affinity for ACE2 (from −50.26 to −56.37 Kcal/mol as compared to −67.49 Kcal/mol of N510Y).

**Fig 6.**
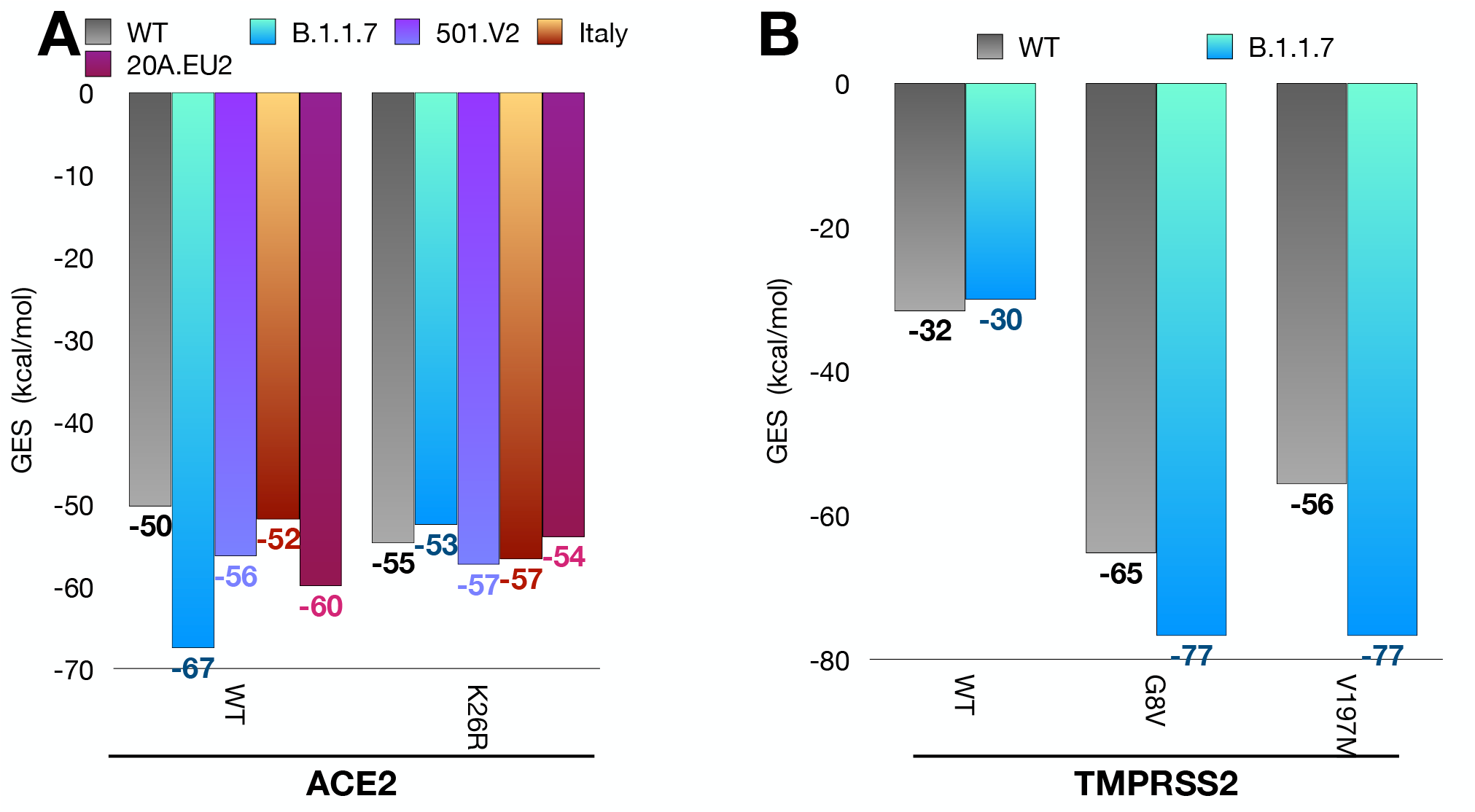
Computed affinity of SARS-CoV-2 variants and wild type or polymorphic variants of ACE2 or TMPRSS2. **A)** Computed affinity of molecular docking complexes between wild type or K26R variant of ACE2, and wild type Spike or emerging variants of Spike with either single (B.1.1.7; 20A.EU2, Italy) or multiple (501.V2) mutations in RBD. **B)** Computed affinity of molecular docking complexes between wild type, G8V or V197M TMPRSS2 variants and emerging variant of Spike B.1.1.7.

Similarly, we decided to evaluate the combined effects of the multiple amino acid substitutions of the SARS-CoV-2 Spike variant B.1.1.7 on computed affinity with TMPRSS2 (Fig. 6B). *De novo* docking simulations were carried out on Gramm-X server, and molecular docking models were screened by FireDock to determine the energy score. Surprisingly, the combined effects were now almost negligible in terms of GES as compared to wild type Spike, while the affinity with either the G8V or V197 TMPRSS2 variants was apparently increased (from −65.30 to −76.77 Kcal/mol, and from −55.74 to −76.77 Kcal/mol, respectively). This result would suggest that higher infectivity of the SARS-CoV-2 B.1.1.7 variant could be mostly due to the N501Y substitution in the Spike RBD-RBM.

## Discussion

In principle, any new infectious agent that challenges a totally susceptible population with little or no immunity against it is able to totally infect the population causing pandemics. Pandemics rapidly spread affecting a large part of people causing plenty of deaths with significant social disruption and economic loss. However, if we look at the even worst pandemics in the human history we can realize that ethnic and geographical differences in the susceptibility to disease actually exist, in spite of transmission routes that are the same for all individuals [34]. Infectious sources are susceptible to evolution, and selective pressure by host characteristics and measure to control the pandemic may lead to emergence of more aggressive or indolent strains.

Although with limitations and caveats of *in silico* technology, this study tries to address the question of how some mutations of the Spike protein of SARS-CoV-2 may affect the host-pathogen interactions, providing interesting insight on factors associated with a different individual susceptibility to COVID-19. To alleviate these limitations, we used a combination of bioinformatics tools, and tested different models.

One year after the spread of the SARS COVID-19, its worldwide distribution remains extremely uneven. Lethality is even more inhomogeneous among and within countries. Although differences in mortality might have various causes, including access and efficiency of health systems, total number of people tested, presence and severity of symptoms in tested populations, they are so impressive that it seems legitimate to search for other factors possibly related to individuals as the elements of a population challenged with different types (wild type or mutants) of viruses. Ultimately, infectivity and lethality do not seem linearly related, and probably represent problems to be solved with different, albeit complementary, approaches.

Basic aspects of epidemiology of the disease warrant some considerations: women are probably more prone to infection but often present a less severe disease. Although higher incidence of cardiac, respiratory and metabolic co-morbidities are probably responsible for more severe form of infection in men, estrogen-induced upregulation of ACE2 expression would explain increased susceptibility of women to a less severe and often asymptomatic form of disease. Furthermore, the ACE2 gene is located on Xp22, in an area where genes are reported to escape from X-inactivation, further explaining higher expression in females [35,36].

ACE2 plays an essential role in the renin-angiotensin-aldosterone system, and its loss of function due to the massive binding of viral particles and internalization could constitute an essential element of the pathophysiology of pulmonary and cardiac damage during COVID-19 infection [37,38]. In this context it should be underlined that ACE2 probably plays a dual role in the dynamic of infection and disease course. While at beginning ACE2 overexpression may increase the entry of the virus into the cell and its replication, its consequent viral-induced loss of function results in an unopposed accumulation of angiotensin II that further aggravates the acute lung injury response to viral infection. Indeed, in the rodent blockade of the renin-angiotensin-aldosterone system limits the acute lung injury induced by the SARS-CoV-1 Spike protein [39], suggesting that if ACE2 function is preserved (because of increased baseline expression, as especially seen in pre-menopausal women), clinical course of infection might be less severe.

Amount of ACE2 (whose expression is modulated by different factors, including age and different medical conditions) is only one aspect of the question: it seems clear that the affinity of the virus Spike for ACE2 is a key determinant of its infective potential. In order to choose the experimental model capable of reproducing the essential aspects of human infection, Chan and colleagues [40] determined *in silico* the Spike / ACE2 affinity in primates and in a series of experimental animals, observing that the binding energy is maximal in primates (−62.20 Rosetta energy units (REU)), intermediate in Syrian hamster (−49.96 REU), lower in bat (−39.60 REU). This allowed the authors to predict that hamsters could be infected, which was experimentally confirmed –underlining the reliability of *in silico* modeling- and could be subsequently at the origin of inter-animal transmission. Of note, in the same study, Chan and colleagues [40] showed that the binding energy between ACE2 and Spike of SARS-CoV, responsible for the 2002 epidemic, was −39.49 REU as compared to −58.18 of human ACE2. It has been suggested that polymorphisms in the ACE2 gene could reduce or enhance the wild type Spike affinity with ACE2: *in silico* models have predicted that two non-infrequent polymorphisms −the S19P (0.3% of African populations) and K26R (0.5% of Europeans)-are associated with decreased or increased affinity and their distribution among patients with COVID-19 infection is currently being investigated.

In the present study, we confirmed our previous theoretical hypothesis that N501Y mutation of Spike would be associated with an increased ACE2 affinity, and molecular docking clearly shows that interaction with ACE2 is even better than that of the wild type Spike protein, per se already remarkable. To date, almost all the cases with this mutation have been described in England and Wales (SARS-CoV-2 variant, named B.1.1.7), and in South Africa (SARS-CoV-2 variant, named 501.V2). The 501.V2 variant also has K417N and E484K in Spike RBM-RBM, in addition to N501Y. Speculations on geographical distribution of these variants and, possibly, interaction with differently distributed ACE2 SNP are not possible. The possible clinical emergence of the N501Y mutation has been also predicted in a recent work by Gu et al [41]: while developing animal models of infection, they adapted a clinical isolate of SARS-CoV-2 by serial passaging in the respiratory tract of aged BALB/c mice. At passage 6 the resulting mouse-adapted strain showed increased infectivity and a pathology similar to severe human disease (interstitial pneumonia) in both young and aged mice after intranasal inoculation. Deep sequencing revealed a panel of adaptive mutations, including the N501Y mutation that is located at the RBD of the Spike protein. By using a different molecular docking approach, the authors showed an increased ACE2/Spike affinity associated with this mutation. In the present study, we showed a substantial increase in computed binding affinity with a global energy score rising from −50.26 Kcal/mol in the wild type Spike to −67.49 Kcal/mol in the N501Y Spike: one should consider that biological effects are probably much more important than mere physical variation in global energy score. Of note when measuring the affinity of N501Y Spike for K26R ACE2 variant (which has a significantly higher affinity for wild type Spike as compared to wild type ACE2), we observed a decrease in affinity (from −54.79 Kcal/mol to −52.57), suggesting that the effect of N501Y substitution is dependent on ACE2 genetic background. We provided theoretical evidence through the analysis of molecular docking complexes by Chimera that the number and the arrangement of contacts and H-bonds were different in the different complexes, with a higher contact density in N501Y Spike RBD / wild type ACE2 explaining enhanced affinity.

In our theoretical approach, the effect of the other leading mutation of Spike, namely D614G, may occur through interaction with TMPRSS2. This mutation involves a region far from the RBD and *in vitro* studies showed that neutralization by monoclonal antibodies targeting the RBD has equal effects when either D614 or D614G are challenged. Our models predict that D614G replacement drastically change the peptide secondary structure by replacing the C-terminal β-sheet with a α-helix in the region close to the mutation, and our CABSflex simulations show an increase in flexibility of the region of interest. Thus, changes in whole protein conformation could results in variations in affinity between ACE2 and D614G Spike. In the recent work by Yurkovetskiy et al. [42], who also showed D614G changes in the conformation of the S1 domain in the SARS-CoV-2 S by a cryo-electron microscopy approach, the association rate between D614G and ACE2 was slower than that between D614 and ACE2, and the dissociation rate of D614G was faster, resulting in a lower affinity. As the mutation occurs in the region of the Spike interacting with TMPRSS2, in our approach, we focused on the possible impact of these possibly modified interactions to explain global changes in interactions between virus and host machineries. Thus, we observed that D614G change resulted in a notable increase in the computed affinity of SARS-CoV-2 Spike protein for wild type TMPRSS2. Chimera modeling enlightened molecular interactions between the wild type or mutant Spike and TMPRSS2 and showed how changes in H-bond was at the origin of the increase in affinity. We also modeled the interactions between either D614 or D614G with TMPRSS2, in both its wild type and polymorphic forms. Of note polymorphisms of TMPRSS2 are frequent, accounting sometimes for 10 percent of populations. These models showed that polymorphic forms of TMPRSS2 are associated with significant changes in the binding of Spike with either its wild type form or the D614G variant, providing a possible further explanation for differences in the diffusion rates of the infection.

These elements prompt us to conclude that if interactions between wild type Spike, ACE2 and TMPRSS2 have been identified as the basic mechanism of COVID-19 infection at cellular level, polymorphic variation of not only Spike but also host’s protein are likely to be determinant of the possibility of infection, and, may be, clinical course. Another point of interest is the possible effect of multiple Spike mutations. This point has to do with the evolution of the virus in the human host. The D614G is also present in the SARS-CoV-2 variant B.1.1.7, together with additional amino acid substitutions A570D, P681H, and T716I mapping in the region between the Fusion Peptide and the RBD domain. Intriguingly, while these single amino acid replacements increases the computed affinity for wild type TMPRSS2, and results in different affinities for G8V and V197M TMPRSS2 variants depending on specific substitutions, their combination in the B.1.1.7 Spike variant does not seem to affect substantially the affinity of the mutated Spike for wild type TMPRSS2, while it seems to increase the affinity for TMPRSS2 variants. This result would suggest, on one hand, that higher infectivity of the SARS-CoV-2 B.1.1.7 variant could be mostly due to the N501Y substitution in the Spike RBD-RBM, and, secondly, that the effects of multiple Spike mutations are mostly dependent on the host genetic background.

## Materials and methods

### Dataset analysis

Frequencies of SARS-CoV-2 Spike protein variants were previously reported [24,25]. Dataset S1 of Supplementary material [24] contains more than 32.000 sequences of SARS-CoV-2 Spike; Dataset S2 of Supplementary material [25] contains the relative frequency of each SARS-CoV-2 Spike missense. Dataset S3 of Supplementary material includes 29 sequences of SARS-CoV previously reported [26,27]The functional domains of protein were mapped by CD-search [43] while the trans-membrane and inner domains were predicted by TMHMM Server v. 2.0. [44] and confirmed by UniProtKB (ID: P0DTC2 SPIKE_SARS2). UniProtKB also provides information about glycosylation and disulfide bond sites. Datasets S1-3 were used to determine the relative distribution of SARS-CoV and SARS-CoV-2 Spike variants with respect to each protein domain. Datasets S1 was used to assess the geographical distribution and the temporal spread of the SARS-CoV-2 Spike D614G variant. Dataset S4 of Supplementary material was assembled by using Blast-P [45] searching for specific taxonomic groups: Avian coronavirus (taxid:694014), SARS-like coronavirus (taxid:694009), Porcine coronavirus HKU15 (taxid:1159905), Bovine coronavirus (taxid:11128) and Bat coronavirus (taxid:1508220). NCBI reference sequence of SARS-CoV-2 spike protein was used as query sequence (YP_009724390.1). ClustalO mutli-alignment tool [46] was used to identify, in the Spike proteins of examined coronaviruses, the amino acid residue corresponding to the amino acid residue 614 of SARS-CoV-2 Spike. GnomAD database v2.1.1 was used to gain information about the TMPRSS2 polymorphisms, and frequencies in human populations. This set of data was also analyzed by multivariate methods (NM-MDS and PCA) using PAST [47].

### Structural dynamical feature analyses

Secondary structures of the region spanning the amino acids 601-627 in the SARS-CoV-2 Spike protein (GTNTSNQVAVLYQDVNCTEVPVAIHAD), and the corresponding region (amino acids 587-613) of SARS-CoV Spike (GTNTSSEVAVLYQDVNCTDVSTAIHAD) was predicted by using the *ab initio* method on PEPfold server [29,30]. The same analysis was performed with SARS-CoV-2 Spike and SARS-CoV Spike protein variants (D614E, D614G, D614A, D614P in SARS-CoV-2 Spike and the corresponding variants in SARS-CoV Spike). CABSflex [31] was used to analyze the possible effects of the amino acid substitutions on flexibility of these peptides. All images were processed and visualized by Chimera USCF [48].

### Molecular docking simulations

The model of isoform 1 of TMPRSS was obtained by using I-Tasser server [49] with the sequence NP_001128571.1 as input. The model of the trimeric SARS-CoV-2 Spike was downloaded from Zhang Laboratory web page (https://zhanglab.ccmb.med.umich.edu/COVID-19/). The docking simulations were carried out on Gramm-X server [32] using the chain A of Spike as a receptor and TMPRSS2 as a ligand. The molecular docking models were screened by FireDock to determine the energy score [50,51]. The SSIPe server [52,53] was used to obtain the models of the protein variants starting with the wild type Spike and TMPRSS2 proteins. These models were re-docked by using Gramm-X [32]. Using Chimera [49], the obtained complexes were analyzed to select those that show an interaction between TMPRSS2 and the Spike domain of cleavage sites. The final complexes were analyzed by using FindHbond and FindClashes/Contacts tools of Chimera in order to identify contacts, pseudobonds and H-bonds. FindClashes/Contacts is a tool that allows the identification of interatomic contacts using van der Waals (VDW) radii. This method was used to localize all direct (polar and nonpolar) interactions between two atoms, including both unfavorable (clashes) and favorable interactions.

In order to compare the Global Energy Scores (GES) in the interaction between wild type or N501Y Spike RBD with wild type or K26R ACE2 the HDOCK server was used [54]. This tool was based on a hybrid method: template-base modeling and *ab initio* docking. Template based modeling was choose as the best strategy to compute the affinity between RBD and ACE2 because many ACE2-RBD crystallographic models were reported in public database. On the other hand, Gramm-x [32], a tool based on rigid-body docking (Lennard-Jones modified function), was used to identify novel binding sites. Using the HDOCK server [54], receptors and ligands were submitted as primary amino acid sequences, including wild type and variant sequences (K26R ACE2 and N501Y RBD). Docking complexes obtained with the template-base modeling were selected, and the affinity energy was calculated using FireDock [50,51].

The workflow used to perform docking simulations was reported in Fig. S5.

## Conflict of interest

The authors declare no conflict of interest.

## Authors’ contribution

M.C. contributed to experimental set-up, pipeline development, *in silico* analysis; P.F. contributed to study designing and data providing; M.A. and P.A. contributed to coordination, conception, designing and writing. M.A. and P.A. contributed equally to the work. All authors critically revised draft versions of the manuscript and approved the final version.

## Acknowledgements

The work was partially supported by a grant from “Fondation du Souffle, Cohortes en Pneumologie 2020”

The authors are grateful to Dr Niel Insdorf for critical relecture of the paper.

## Additional information

Supplementary Figures and Supplementary Tables can be found in the Supplementary Material section of the online article.

## Supplementary material

**Fig S1.**
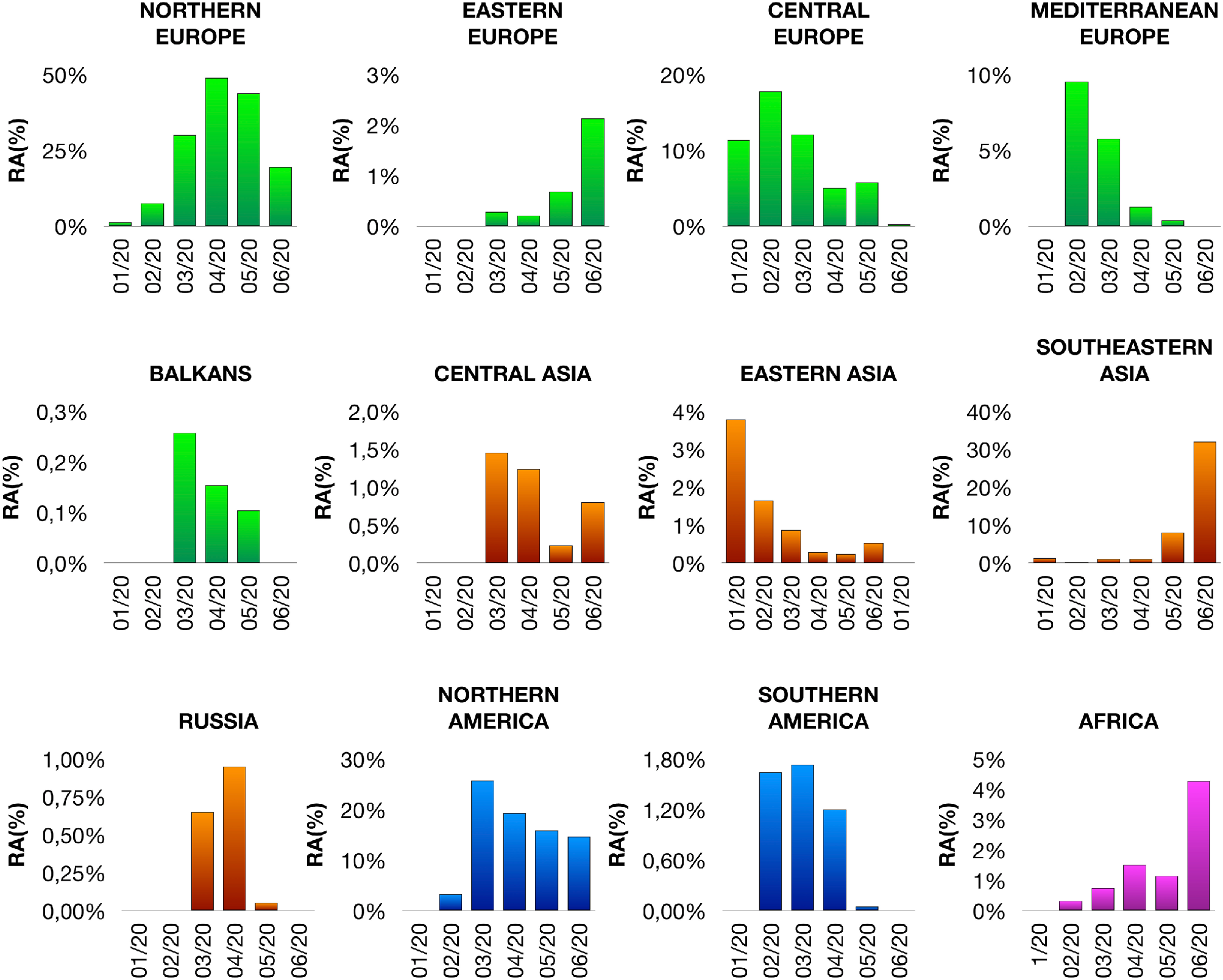
Geographical distribution and temporal spread of the SARS-CoV-2 Spike D614G variant. Geographical distribution and temporal spread of the SARS-CoV-2 Spike D614G variant (time range: 01/2020-06/2020) in 5 regions of Europe (green), 4 regions of Asia (orange), 2 regions of America (blue) and Africa (magenta) are shown. RA, relative abundance.

**Fig S2.**
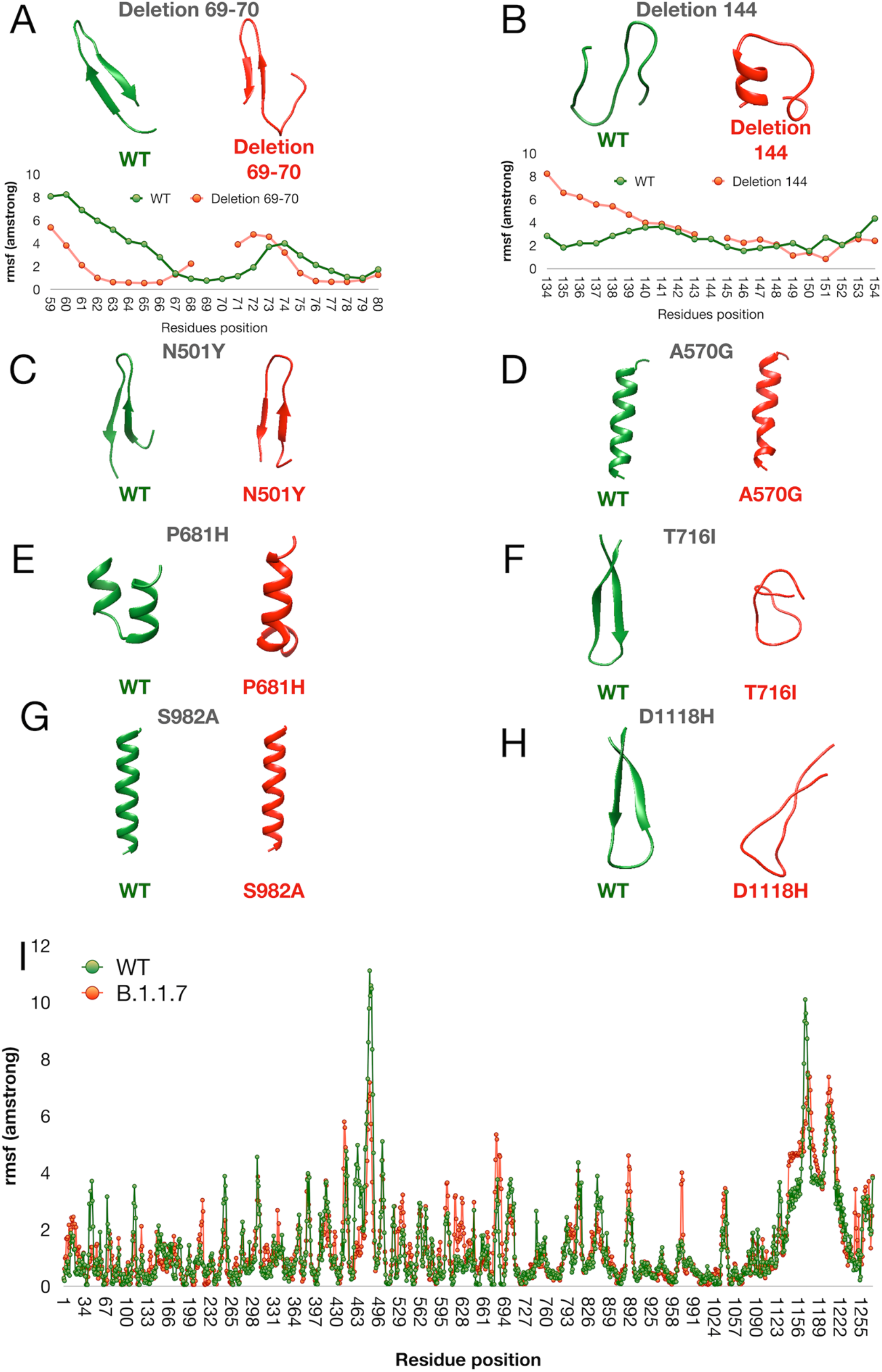
Regional secondary structures prediction and domain flexibility of wild type and B.1.1.7 variants of Spike of SARS-CoV-2. **A and B)** Effects of two N-terminal deletions (69-70 and 144) on the secondary structure and flexibility. **C-G)** Effects of amino acid substitutions on local secondary structure of Spike. **I)** Comparison of flexibility between wild type and B.1.1.7 Spike.

**Fig S3.**
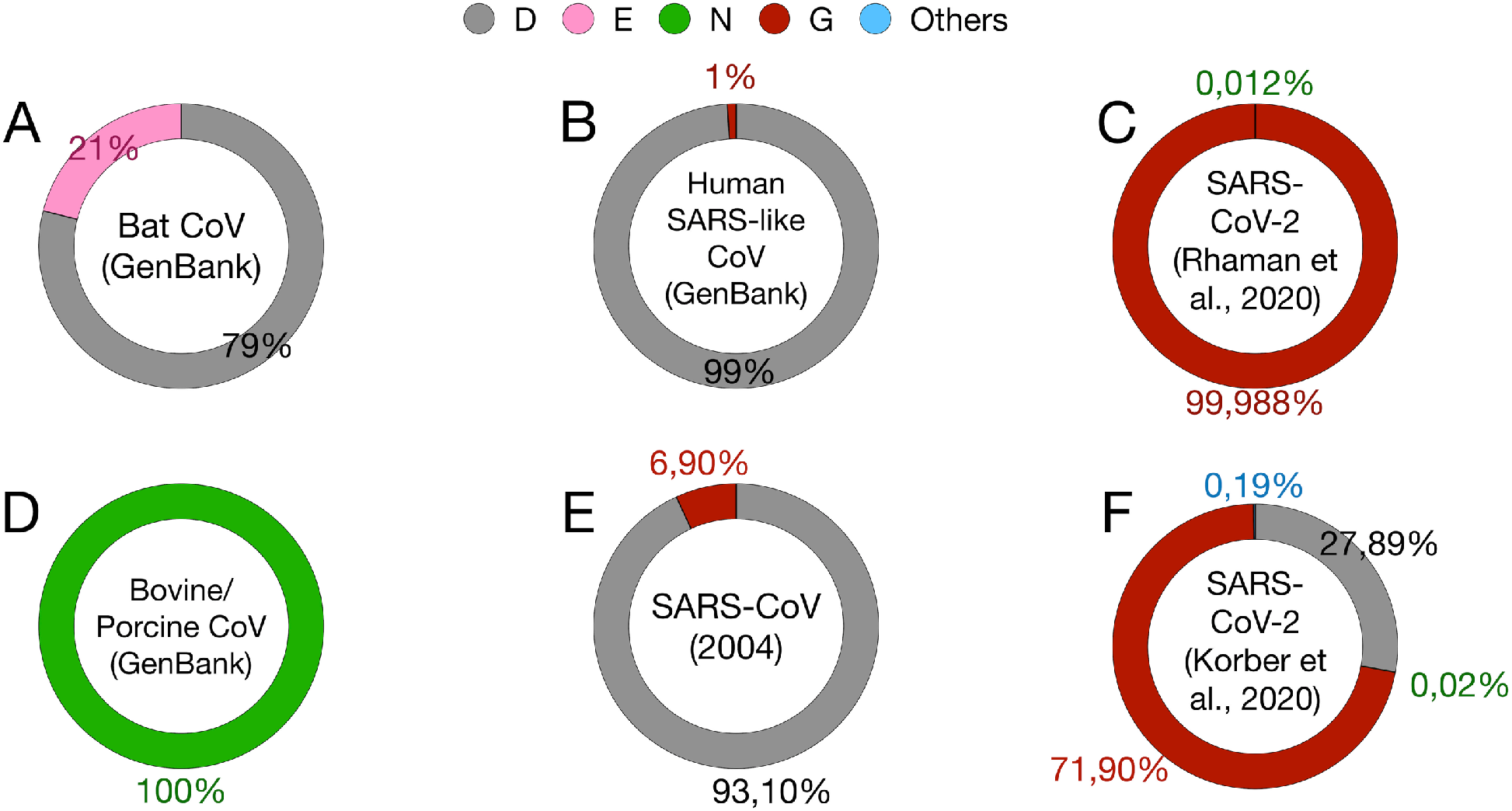
Frequencies of the most represented amino acids at the location 614 in SARS-like viruses. Frequencies reported from the diverse datasets: SARS-CoV-2 (Dataset S1, Dataset S2); SARS-CoV-1 (Dataset S3); SARS-like human virus (Dataset S4); Bat, Avian, Bovine and Porcine Coronavirus (Dataset S4).

**Fig S4.**
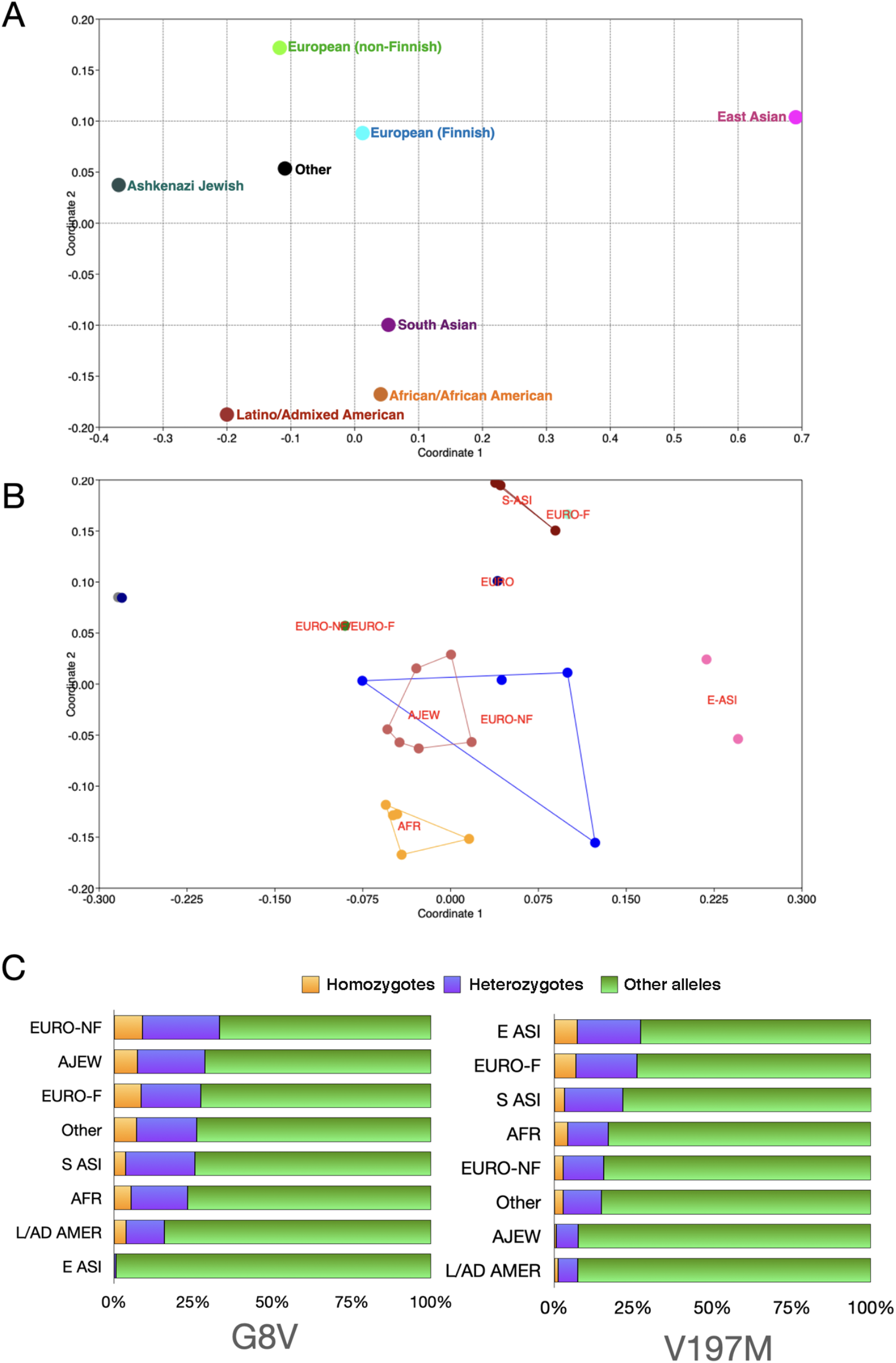
Frequencies of TMPRSS2 polymorphisms among human population. **A**) Ordination plot (NM-MDS) that compare all TMPRSS2 variants in 7 human populations. **B)** Ordination plot (PCA) distribution of most diffused TMPRSS2 variants (relative frequencies >0.05) in 7 human populations. **C)** Frequencies of two TMPRSS2 variants (G8V and V197M) in homozygous or heterozygous conditions. Frequencies other alleles were determined as described in materials and Methods. AFR, African/African-American; AMR, Latino/Admixed American; ASJ, Ashkenazi Jewish; EAS, East Asian; FIN, Finnish; NFE, Non-Finnish European; SAS, South Asian.

**Fig S5.**
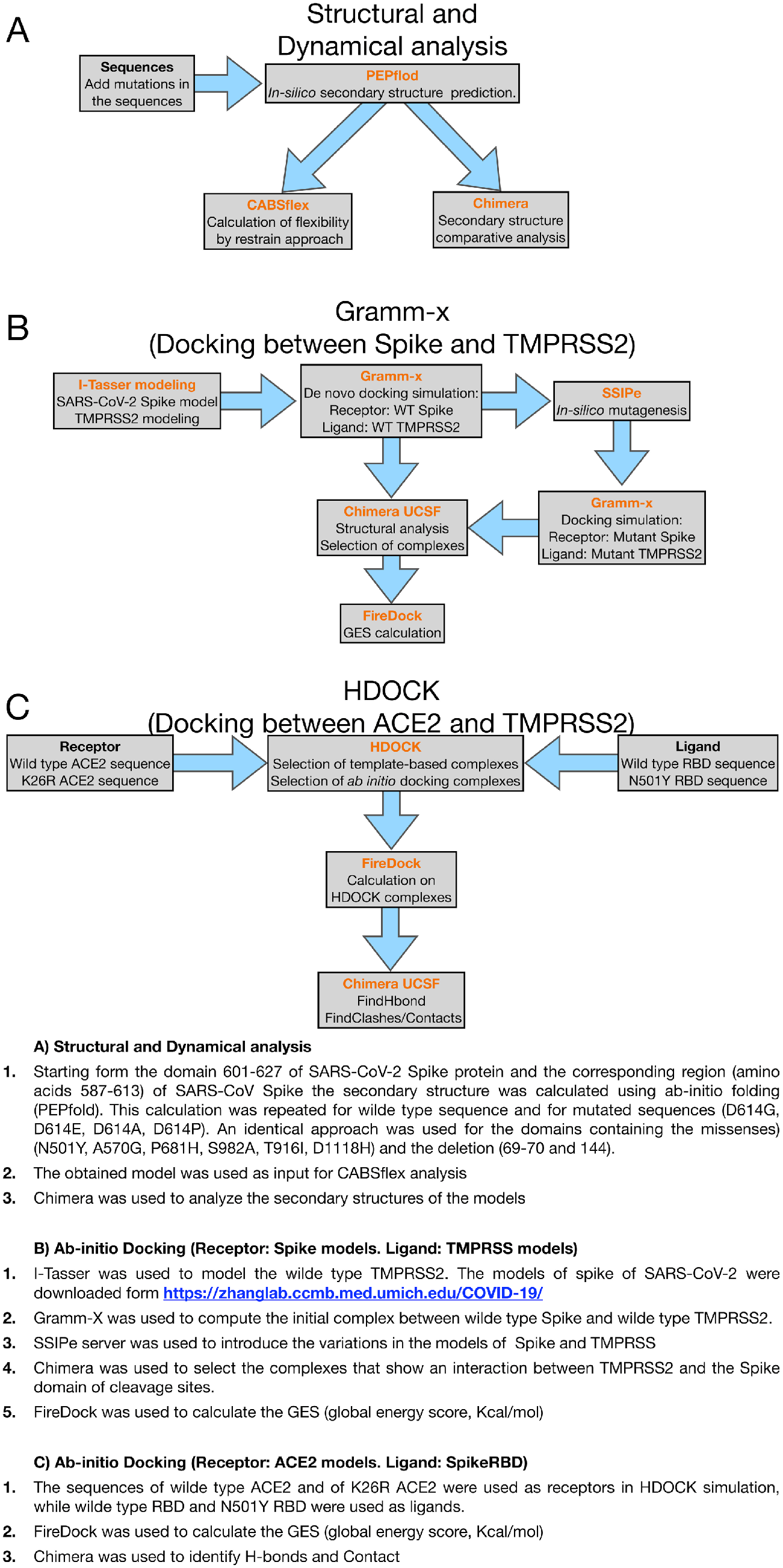
Computational workflow used in this study. **A)** Structural and dynamical analysis performed on 601-627 domain of SARS-CoV-2 Spike protein and the corresponding region (587-613) of SARS-CoV Spike. This analysis was repeated for 4 missenses (D614G, D614E, D614A, D614P). An identical approach was used for the domains containing the missenses (N501Y, A570G, P681H, S982A, T916I, D1118H) and the deletions (69-70 and 144). **B)** Ab-initio docking was used to characterize the interaction between WT or variant Spike proteins and WT or variant TMPRSS2. **C)** Template base docking used to compute the effect of RBD N501Y missense on the interaction with WT or K26R ACE2.

## Supplementary Datasets

**Dataset S1**. The Dataset reports the missense mutations identified in the SARS-CoV-2 Spike proteins described by Rahman et al. [24]. **Table 1**. Counts and frequencies of identified missense mutations. **Table 2**. Temporal distribution of missense mutations. **Table 3**. Geographical distribution of missense mutations. **Table 4:** Countries included in the geographical region. **Table 5**. Distribution of the amino acid variations in the Spike domain.

**Dataset S2**. The Dataset reports the frequencies of the missense mutations identified in the SARS-CoV-2 Spike proteins described by Kromer et al. [25].

**Dataset S3**. The Dataset reports the missense mutations identified in two studies on SARS-CoV in the year 2004 [26,27]. **Table 1**. Counts and frequencies of identified missense mutations. **Table 5**. Distribution of the amino acid variations in the Spike domain.

**Dataset S4**. The Dataset was implemented using BLAST, restricting the search to: i) the SARS-CoV-2 sequences, SARS-Like human virus, Bat SARS-Like virus and porcine/bovine SARS-like virus. Ref sequences of SARS-CoV-2 was used as query.

## Notes

### Competing Interest Statement

The authors have declared no competing interest.

