## Supplemental Datasets 4 for "The lethal triad: SARS-CoV-2 Spike, ACE2 and TMPRSS2. Mutations in host and pathogen may affect the course of pandemic"

```
Avian

REF                          EILDITPCSFGGVSVITPGTNTSNQVAVLYQDVNCTEVPVAIHADQLTPTWRVYSTGSNV	642
YP_009825008.1:366-1140      SLYTVKPCSTVSTQAVIVA----NQLAGLYLPLSCDIVFNLNLGNETTPVDGGCLVFNST	159
ADP06481.1:429-1094          SYYKVNPCSDINEQYVVSG----GNLVGKLTSNNQTVAQQL--------GDMFYVKFSTS	100
QDM39239.1:422-1115          HYYKINPCNDVNQQYVVSG----GNIVGLLTSSNETGSTQL--------EDQFYIKLTNS	107
QKV27925.1:423-1116          HYYKINPCNDVNQQYVVSG----GNIVGLLTSSNETGSIQL--------EDQFYIKLTNS	107
AJP08845.1:400-1120          NYYKVNPCKDVNQQYVVSG----GNIVGLLTSINSSGSQLL--------EDQYYVRLTNT	141


BAT
REF                     VITPGTNTSNQVAVLYQDVNCTEVPVAIHADQLTPTWRVYSTGSNVFQTRAGCLIGAEHV	656
AGZ48806.1:3-1256       VITPGTNTSSEVAVLYQDVNCTDVPVAIHADQLTPSWRVYSTGNNVFQTQAGCLIGAEHV	641
ATO98132.1:3-1256       VITPGTNTSSEVAVLYQDVNCTDVPVAIHADQLTPSWRVYSTGNNVFQTQAGCLIGAEHV	641
ATO98218.1:3-1256       VITPGTNTSSEVAVLYQDVNCTDVPVAIHADQLTPAWRIYSTGNNVFQTQAGCLIGAEHV	641
ATO98231.1:3-1256       VITPGTNTSSEVAVLYQDVNCTDVPVAIHADQLTPAWRIYSTGNNVFQTQAGCLIGAEHV	641
AGZ48828.1:3-1256       VITPGTNTSSEVAVLYQDVNCTDVPVAIHADQLTPSWRVHSTGNNVFQTQAGCLIGAEHV	641
AGZ48818.1:3-1256       VITPGTNTSSEVAVLYQDVNCTDVPVAIHADQLTPSWRVYSTGNNVFQTQAGCLIGAEHV	641
ATO98157.1:1-1255       VITPGTNTSSEVAVLYQDVNCTDVPVAIHADQLTPAWRIYSTGNNVFQTQAGCLIGAEHV	642
ATO98205.1:1-1255       VITPGTNTSSEVAVLYQDVNCTDVPVAIHADQLTPSWRVYSTGNNVFQTQAGCLIGAEHV	642
AAV97990.1:1-1255       VITPGTNASSEVAVLYQDVNCTDVSTLIHADQLTPAWRIYSTGNNVFQTQAGCLIGAEHV	642
AAV98000.1:1-1255       VITPGTNASSEVAVLYQDVNCTDVSTLIHAEQLTPAWRIYSTGNNVFQTQAGCLIGAEHV	642
AAU04662.1:1-1255       VITPGTNASSEVAVLYQDVNCTDVSTLIHAEQLTPAWRIYSTGNNVFQTQAGCLIGAEHV	642
AAV91631.1:1-1255       VITPGTNASSEVAVLYQDVNCTDVSTLIHAEQLTPAWRIYSTGNNVFQTQAGCLIGAEHV	642
AAU04649.1:1-1255       VITPGTNASSEVAVLYQDVNCTDVSTLIHAEQLTPAWRIYSTGNNVFQTQAGCLIGAEHV	642
AAU04646.1:1-1255       VITPGTNASSEVAVLYQDVNCTDVSTLIHAEQLTPAWRIYSTGNNVFQTQAGCLIGAEHV	642
AAV98001.1:1-1255       VITPGTNASSEVAVLYQDVNCTDVSTLIHAEQLTPAWRIYSTGNNVFQTLAGCLIGAEHV	642
AAV97998.1:1-1255       VITPGTNASSEVAVLYQDVNCTDVSTLIHAEQLTPAWRIYSTGNNVFQTQAGCLIGAEHV	642
AAV49720.1:1-1255       VITPGTNASSEVAVLYQDVNCTDVSTLIHADQLTPAWRIYSTGNNVFQTQAGCLIGAEHV	642
AAV97985.1:1-1255       VITPGTNASSEVAVLYQDVNCTDVSTLIHAEQLTPAWRIYSTGNNVFQTQAGCLIGAEHV	642
AAU93319.1:1-1255       VITPGTNASSEVAVLYQDVNCTDVSTLIHAEQLTPAWRIYSTGNNVFQTQAGCLIGAEHV	642
AAV97992.1:1-1255       VITPGTNASSEVAVLYQDVNCTDVSTLIHAEQLTPAWRIYSTGNNVFQTQAGCLIGAEHV	642
AAV49722.1:1-1255       VITPGTNASSEVAVLYQDVNCTDVSTLIHAEQLTPAWRIYSTGNNVFQTQAGCLIGAEHV	642
AAU04664.1:1-1255       VITPGTNASSEVAVLYQDVNCTDVSTLIHAEQLTPAWRIYSTGNNVFQTQAGCLIGAEHV	642
AAV97989.1:1-1255       VITPGTNASSEVAVLYQDVNCTDVSTLIHAEQLTPAWRIYSTGNNVFQTQAGCLIGAEHV	642
AAS10463.1:1-1255       VITPGTNASSEVAVLYQDVNCTDVSTLIHAEQLTPAWRIYSTGNNVFQTQAGCLIGAEHV	642
AAV49723.1:1-1255       VITPGTNASSEVAVLYQDVNCTDVSTLIHAEQLTPAWRIYSTGNNVFQTQAGCLIGAEHV	642
AAV49719.1:1-1255       VITPGTNASSEVAVLYQDVNCTDVSTLIHAEQLTPAWRIYSTGNNVFQTQAGCLIGAEHV	642
AAV97986.1:1-1255       VITPGTNASSEVAVLYQDVNCTDVSTVIHAEQLTPAWRIYSTGNNVFQTQAGCLIGAEHV	642
AAV97984.1:1-1255       VITPGTNASSEVAVLYQDVNCTDVSTLIHAEQLTPAWRIYSTGNNVFQTQAGCLIGAEHV	642
AAV98002.1:1-1255       VITPGTNASSEVAVLYQDVNCTDVSTLIHAEQLTPAWRIYSTGNNVFQTQAGCLIGAEHV	642
AAU04661.1:1-1255       VITPGTNASSEVAVLYQDVNCTDVSTLIHAEQLTPAWRIYSTGNNVFQTQAGCLIGAEHV	642
AAV97995.1:1-1255       VITPGTNASSEVAVLYQDVNCTDVSTLIHAEQLTPAWRIYSTGNNVFQTQAGCLIGAEHV	642
AAS00003.1:1-1255       VITPGTNASSEVAVLYQDVNCTDVSTAIHADQLTPAWRIYSTGNNVFQTQAGCLIGAEHV	642
AAP51227.1:1-1255       VITPGTNASSEVAVLYQDVNCTDVSTAIHADQLTPAWRIYSTGNNVFQTQAGCLIGAEHV	642
AAP82968.1:1-1255       VITPGTNASSEVAVLYQDVNCTDVSTAIHADQLTPAWRIYSTGNNVFQTQAGCLIGAEHV	642
ABF68955.1:1-1255       VITPGTNASSEVAVLYQDVNCTDVSTAIHADQLTPAWRIYSTGNNVFQTQAGCLIGAEHV	642
AAR07631.1:1-1255       VITPGTNASSEVAVLYQDVNCTDVSTAIHADQLTPAWRIYSTGNNVFQTQAGCLIGAEHV	642
AAR07628.1:1-1255       VITPGTNASSEVAVLYQDVNCTDVSTAIHADQLTPAWRIYSTGNNVFQTQAGCLIGAEHV	642
AAR07627.1:1-1255       VITPGTNASSEVAVLYQDVNCTDVSTAIHADQLTPAWRIYSTGNNVFQTQAGCLIGAEHV	642
AAR07626.1:1-1255       VITPGTNASSEVAVLYQDVNCTDVSTAIHADQLTPAWRIYSTGNNVFQTQAGCLIGAEHV	642
AAR07629.1:1-1255       VITPGTNASSEVAVLYQDVNCTDVSTAIHADQLTPAWRIYSTGNNVFQTQAGCLIGAEHV	642
AAR07625.1:1-1255       VITPGTNASSEVAVLYQDVNCTDVSTAIHADQLTPAWRIYSTGNNVFQTQAGCLIGAEHV	642
ABF68958.1:1-1255       VITPGTNASSEVAVLYQDVNCTDVSTAIHADQLTPAWRIYSTGNNVFQTQAGCLIGAEHV	642
AAT74874.1:1-1255       VITPGTNASSEVAVLYQDVNCTDVSTAIHADQLTPAWRIYSTGNNVFQTQAGCLIGAEHV	642
AAR07624.1:1-1255       VITPGTNASSEVAVLYQDVNCTDVSTAIHADQLTPAWRIYSTGNNVFQTQAGCLIGAEHV	642
AAT76147.1:1-1255       VITPGTNASSEVAVLYQDVNCTDVSTAIHADQLTPAWRIYSTGNNVFQTQAGCLIGAEHV	642
AAR91586.1:1-1255       VITPGTNASSEVAVLYQDVNCTDVSTAIHADQLTPAWRIYSTGNNVFQTQAGCLIGAEHV	642
AAR07630.1:1-1255       VITPGTNASSEVAVLYQDVNCTDVSTAIHADQLTPAWRIYSTGNNVFQTQAGCLIGAEHV	642
AAP13567.1:1-1255       VITPGTNASSEVAVLYQDVNCTDVSTAIHADQLTPAWRIYSTGNNVFQTQAGCLIGAEHV	642
AAX16192.1:1-1255       VITPGTNASSEVAVLYQDVNCTDVSTAIHADQLTPAWRIYSTGNNVFQTQAGCLIGAEHV	642
ACZ71797.1:1-1255       VITPGTNASSEVAVLYQDVNCTDVSTAIHADQLTPAWRIYSTGNNAFQTQAGCLIGAEHV	642
ACZ71961.1:1-1255       VITPGTNASSEVAVLYQDVNCTDVSTAIHADQLTPAWRIYSTGNNVFQTQAGCLIGAEHV	642
ACZ71976.1:1-1255       VITPGTNASSEVAVLYQDVNCTDVSTAIHADQLTPAWRIYSTGNNVFQTQAGCLIGAEHV	642
ACZ72195.1:1-1255       VITPGTNASSEVAVLYQDVNCTDVSTAIHADQLTPAWRIYSTGNNVFQTQAGCLIGAEHV	642
ACZ72254.1:1-1255       VITPGTNASSEVAVLYQDVNCTDVSTAIHADQLTPAWRIYSTGNNVFQTQAGCLIGAEHV	642
ACZ72020.1:1-1255       VITPGTNASSEVAVLYQDVNCTDVSTAIHADQLTPAWRIYSTGNNVFQTQAGCLIGAEHV	642
ACZ71826.1:1-1255       VITPGTNASSEVAVLYQDVNCTDVSTAIHADQLTPAWRIYSTGNNVFQTQAGCLIGAEHV	642
ACZ72108.1:1-1255       VITPGTNASSEVAVLYQDVNCTDVSTAIHADQLTPAWRIYSTGNNVFQTQAGCLIGAEHV	642
ACB69905.1:1-1255       VITPGTNASSEVAVLYQDVNCTDVSTAIHADQLTPAWRIYSTGNNVFQTQAGCLIGAEHV	642
ACB69894.1:1-1255       VITPGTNASSEVAVLYQDVNCTDVSTAIHADQLTPAWRIYSTGNNVFQTQAGCLIGAEHV	642
ACB69883.1:1-1255       VITPGTNASSEVAVLYQDVNCTDVSTAIHADQLTPAWRIYSTGNNVFQTQAGCLIGAEHV	642
BAE93401.1:1-1255       VITPGTNASSEFAVLYQDVNCTDVSTAIHADQLTPAWRIYSTGNNVFQTQAGCLIGAEYV	642
AGT21078.1:1-1255       VITPGTNASSEVAVLYQDVNCTDVSTAIHADQLTPAWRIYSTGNNVFQTQAGCLIGAEYV	642
AFM43867.1:1-1255       VITPGTNASSEVAVLYQDVNCTDVSTAIHADQLTPAWRIYSTGNNVFQTQAGCLIGAEYV	642
ABF68957.1:1-1255       VITPGTNASSEVAVLYQDVNCTDVSTAIHADQLTPAWRIYSTGNNVFQTQAGCLIGAEHV	642
ABF68956.1:1-1255       VITPGTNASSEVAVLYQDVNCTDVSTAIHADQLTPAWRIYSTGNNVFQTQAGCLIGAEHV	642
ABF68959.1:1-1255       VITPGTNASSEVAVLYQDVNCTDVSTAIHADQLTPAWRIYSTGNNVFQTQAGCLIGAEHV	642
BAF42873.1:1-1255       VITPGTNASSEVAVLYQDVNCTDVSTAIHADQLTPAWRIYSTGNNVFQTQAGCLIGAEYV	642
AAR86775.1:1-1255       VITPGTNASSEVAVLYQDVNCTNVSAAIHADQLTPAWRIYSTGNNVFQTQAGCLIGAEHV	642
AEA10443.1:1-1255       VITPGTNASSEVAVLYQDVNCTDVSTAIHADQLTPAWRIYSTGNNVFQTQAGCLIGAEHV	642
ACQ82725.1:1-1255       VITPGTNASSEVAVLYQGVNCTDVSTAIHADQLTPAWRIYSTGNNVFQTQAGCLIGAEHV	642
AFR58714.1:1-1255       VITPGTNASSEVAVLYQDVNCTDVSTAIHADQLTPAWRIYSTGNNVFQTQAGCLIGAEHV	642
ACZ71991.1:1-1255       VITPGTNASSEVAVLYQDVNCTDVSTAIHADQLTPAWRIYSTGNNVFQTQAGCLIGAEHV	642
AEA10473.1:1-1255       VITPGTNASSEVAVLYQDVNCTDVSTAIHADQLTPAWRIYSTGNNVFQTQAGCLIGAEHV	642
ACZ72122.1:1-1255       VITPGTNASSEVAVLYQDVNCTDVSTAIHADQLTPAWRIYSTGNNVFQTQAGCLIGAEHV	642
ABD72995.1:1-1255       VITPGTNASSEVAVLYQDVNCTDVSTAIHADQLTPAWRIYSTGNNVFQTQAGCLIGAEHV	642
ABD72988.1:1-1255       VITPGTNASSEVAVLYQDVNCTDVSTAIHADQLTPAWRIYSTGNNVFQTQAGCLIGAEHV	642
ADC35483.1:1-1255       VITPGTNASSEVAVLYQDVNCTDVSTAIHADQLTPAWRIYSTGNNVFQTQAGCLIGAEHV	642
ABF65836.1:1-1255       VITPGTNASSEVAVLYQDVNCTDVSTAIHADQLTPAWRIYSTGNNVFQTQAGCLIGAEHV	642
AFR58672.1:1-1255       VITPGTNASSEVAVLYQDVNCTDVSTAIHADQLTPAWRIYSTGNNVFQTQAGCLIGAEHV	642
ACZ72093.1:1-1255       VITPGTNASSEVAVLYQDVNCTDVSTAIHADQLTPAWRIYSTGNNVFQTQAGCLIGAEHV	642
AAP33697.1:1-1255       VITPGTNASSEVAVLYQDVNCTDVSTAIHADQLTPAWRIYSTGNNVFQTQAGCLIGAEHV	642
ABD72969.1:1-1255       VITPGTNASSEVAVLYQDVNCTDVSTAIHADQLTPAWRIYSTGNNVFQTQAGCLIGAEHV	642
AAP41037.1:1-1255       VITPGTNASSEVAVLYQDVNCTDVSTAIHADQLTPAWRIYSTGNNVFQTQAGCLIGAEHV	642
ABD72977.1:1-1255       VITPGTNASSEVAVLYQDVNCTDVSTAIHADQLTPAWRIYSTGNNVFQTQAGCLIGAEHV	642
ABD72970.1:1-1255       VITPGTNASSEVAVLYQDVNCTDVSTAIHADQLTPAWRIYSTGNNVFQTQAGCLIGAEHV	642
AAU81608.1:1-1255       VITPGTNASSEVAVLYQDVNCTDVSTAIHADQLTPAWRIYSTGNNVFQTQAGCLIGAEHV	642
ABD73001.1:1-1255       VITPGTNASSEVAVLYQDVNCTDVSTAIHADQLTPAWRIYSTGNNVFQTQAGCLIGAEHV	642
AAR23250.1:1-1255       VITPGTNASSEVAVLYQDVNCTDVSTAIHADQLTPAWRIYSTGNNVFQTQAGCLIGAEHV	642
ABD72972.1:1-1255       VITPGTNASSEVAVLYQDVNCTDVSTAIHADQLTPAWRIYSTGNNVFQTQAGCLIGAEHV	642
ABD72985.1:1-1255       VITPGTNASSEVAVLYQDVNCTDVSTAIHADQLTPAWRIYSTGNNVFQTQAGCLIGAEHV	642
ABD72979.1:1-1255       VITPGTNASSEVAVLYQDVNCTDVSTAIHADQLTPAWRIYSTGNNVFQTQAGCLIGAEHV	642
ABD72984.1:1-1255       VITPGTNASSEVAVLYQDVNCTDVSTAIHADQLTPAWRIYSTGNNVFQTQAGCLIGAEHV	642
P59594.1:1-1255         VITPGTNASSEVAVLYQDVNCTDVSTAIHADQLTPAWRIYSTGNNVFQTQAGCLIGAEHV	642
ABD72982.1:28-1255      VITPGTNASSEVAVLYQDVNCTDVSTAIHADQLTPAWRIYSTGNNVFQTQAGCLIGAEHV	615
QND76020.1:5-1259       VITPGTNASSEVAVLYQDVNCTDVPTAIRADQLTPAWRVYSTGVNVFQTQAGCLIGAEHV	642
QND76034.1:5-1259       VITPGTNASSEVAVLYQDVNCTDVPTAIRADQLTPAWRVYSTGVNVFQTQAGCLIGAEHV	642
AVP78042.1:12-1245      VITPGTNTSSEVAVLYQDVNCTDVPTTIHADQLTPAWRIYAIGTSVFQTQAGCLIGAEHV	621
AVP78031.1:12-1246      VITPGTNTSLEVAVLYQDVNCTDVPTTIHADQLTPAWRIYATGTNVFQTQAGCLIGAEHV	622
QHR63300.2:1-1269       VITPGTNASNQVAVLYQDVNCTEVPVAIHADQLTPTWRVYSTGSNVFQTRAGCLIGAEHV	656
6ZGF_A:32-1252          VITPGTNTSNQVAVLYQDVNCTEVPVAIHADQLTPTWRVYSTGSNVFQTRAGCLIGAEHV	656
                        *******:* :.*****.****:* . *:*:****:**::: * ..*** ********:*


Bovine coronavirus
Porcine coronavirus HKU15
Bovine enteritic coronavirus
REF                       PGTNTSNQVAVLYQDVNCTEVPVAIHADQLTPTWRVYSTGSNVFQTRAGCLIGAEHVNN-	658
ACB30203.1:621-1358       H--ANSSEPALLFRNIKCNYVFNNALSRQLQP--------INYFDSYLGCVVNADNSTSS	123
AVI15053.1:621-1358       H--ANSSEPALLFRNIKCNYVFNNTLSRQLQP--------INYFDSYLGCVVNADNSTSS	123
AMQ23489.1:621-1358       H--ANSSEPALLFRNIKCNYVFNNTLSRQLQP--------INYFDSYLGCVVNADNSTSS	123
AVI15052.1:621-1358       H--ANSSEPALLFRNIKCNYVFNNILSRQLQP--------INYFDSYLGCVVNADNSTSS	123
AMQ23499.1:621-1358       H--ANSSEPALLFRNIKCNYVFNNTLSRQLQP--------INYFDSYLGCVVNADNSTSS	123
AVI15011.1:622-1359       H--ANSSEPALLFRNIKCNYVFNNTLSRQLQP--------INYFDSYLGCVVNADNSTSS	123
AMQ23498.1:621-1358       H--ANSSEPALLFRNIKCNYVFNNTLSRQLQP--------INYFDSYLGCVVNADNSTSS	123
AMQ23497.1:621-1358       H--ANSSEPALLFRNIKCNYVFNNTLSRQLQP--------INYFDSYLGCVVNADNSTSS	123
AMQ23494.1:621-1358       H--ANSSEPALLFRNIKCNYVFNNTLSRQLQP--------INYFDSYLGCVVNADNSTSS	123
AMQ23493.1:621-1358       H--ANSSEPALLFRNIKCNYVFNNTLSRQLQP--------INYFDSYLGCVVNADNSTSS	123
AVI15000.1:621-1358       H--ANSSEPALLFRNIKCNYVFNNTLSRQLQP--------INYFDSYLGCVVNADNSTSS	123
ACT11030.1:621-1358       H--ANSSEPALLFRNIKCNYVFNNTLSRQLQP--------INYFDSYLGCVVNADNSTSS	123
AGO98859.1:621-1358       H--ANSSEPALLFRNIKCNYVFNNTLSRQLQP--------INYFDSYLGCVVNADNSTSS	123
QGW57585.1:621-1358       H--ANSSEPALLFRNIKCNYVFNNTLSRQLQP--------INYFDSYLGCVVNADNSTSS	123
AGO98868.1:621-1358       H--ANSSEPALLFRNIKCNYVFNNTLLRQLQP--------INYFDSYLGCVVNADNSTSS	123
AGO98867.1:621-1358       H--ANSSEPALLFRNIKCNYVFNNTLLRQLQP--------INYFDSYLGCVVNADNSTSS	123
ABP73656.1:621-1358       H--ANSSKPALLFQNIKCNYVFNNTLSRQLQP--------INYFDSYLGCVVNADNSTSS	123
AGO98860.1:621-1358       H--ANSSEPALLFRNIKCNYVFNNTLSRQLQP--------INYFDSYLGCVVNADNSTSS	123
AGO98881.1:621-1358       H--ANSSEPALLFRNIKCNYVFNNTLSRQLQP--------INYFDSYLGCVVNADNSTSS	123
AGO98882.1:621-1358       H--ANSSEPALLFRNIKCNYVFNNTLSRQLQP--------INYFDSYLGCVVNADNSTSS	123
AGO98883.1:621-1358       H--ANSSEPALLFRNIKCNYVFNNTLSRQLQP--------INYFDSYLGCVVNADNSTSS	123
AGO98871.1:621-1358       H--ANSSEPALLFRNIKCNYVFNNTLSRQLQP--------INYFDSYLGCVVNADNSTSS	123
AGO98885.1:621-1358       H--ANSSEPALLFRNIKCNYVFNNTLSRQLQP--------INYFDSYLGCVVNADNSTSS	123
ABD18928.1:621-1358       H--ANSSEPALLFRNIKCNYVFNNTLPRQLQP--------INYFDSYLGCVVNADNSTSS	123
ABD18908.1:621-1358       H--ANSSEPALLFRNIKCNYVFNNTLSRQLQP--------INYFVSYLGCVVNADNSTSS	123
AAX38495.1:621-1358       H--ANSSEPALLFRNIKCNYVFNNTLPRQLQP--------INYFDSYLGCVVIADNSTSS	123
ACG75895.1:621-1358       H--ANSSEPALLFRNIKCNYVFNNTFSRQLQP--------INYFDSYLGCVVNADNSTSS	123
ACB30200.1:621-1358       H--ANSSEPALLFRNIKCNYVFNNTFSRQLQP--------INYFDSYLGCVVNADNSTSS	123
ABD18930.1:621-1358       H--ANSSEPALLFRNIKCNYVFNNTLSRQLQP--------INYFDSYLGCVVNADNSTSS	123
ABD18917.1:621-1358       H--ANSSEPALLFRNIKCNYVFNNTLSRQLQP--------INYFDSYLGCVVNADNSTSS	123
ADP21335.1:621-1358       H--ANSSEPALLFRNIKCNYVFNNTLSRQLQP--------INYFDSYLGCVVNADNSTSS	123
ADP21336.1:621-1358       H--ANSSEPALLFRNIKCNYVFNNTLSRQLQP--------INYFDSYLGCVVNADNSTSS	123
ABD18909.1:621-1358       H--ANSSEPALLFRNIKCNYVFNNTLSRQLQP--------INYFDSYLGCVVNADNSTSS	123
ABD18913.1:621-1358       H--ANSSEPALLFRNIKCNYVFNNTLSRQLQP--------INYFDSYLGCVVNADNSTSS	123
BBM61107.1:621-1358       H--ANSSEPALLFRNIKCNYVFNNTLSRQLQP--------INYFDSYLGCVVNADNSTSS	123
BBM60917.1:621-1358       H--ANSSEPALLFRNIKCNYVFNNTLSRQLQP--------INYFDSYLGCVVNADNSTSS	123
BBM61407.1:619-1356       H--ANSSEPALLFRNIKCNYVFNNTLSRQLQP--------INYFDSYLGCVVNADNSTSS	123
BBM61447.1:621-1358       H--ANSSEPALLFRNIKCNYVFNNILSRQLQP--------INYFDSYLGCVVNADNSTSS	123
BBM60977.1:621-1358       H--SNSSEPALLFRNIKCNYVFNNILSRQLQP--------INYFDSYLGCVVNADNSTSS	123
BBM60967.1:621-1358       H--SNSSEPALLFRNIKCNYVFNSTLSRQLQP--------INYFDSYLGCVVNADNSTSS	123
BBM61057.1:621-1358       H--ANSSEPALLFRNIKCNYVFNNTLLRQLQP--------INYFDSYLGCVVNADNSTSS	123
ABP38243.1:621-1358       H--ANSSEPALLFRNIKCNYVFNNTLSRQLQP--------INYFDSYLGCVVNADNSTSS	123
BBM61387.1:621-1358       H--ANSSEPALLFRNIKCNYVFNNTLSRQLQP--------INYFDSYLGCVVNADNSTSS	123
BBM61377.1:621-1358       H--ANSSEPALLFRNIKCNYVFNNTLSRQLQP--------INYFDSYLGCVVNADNSTSS	123
ACJ66977.1:621-1358       H--ANSSEPALLFRNIKCNYVFNNTLSRQLQP--------INYFDSYLGCVVNADNSTSS	123
ABP38253.1:621-1358       H--ANSSEPALLFRNIKCNYVFNNTLSRQLQP--------INYFDSYLGCVVNADNSTSS	123
BBM61257.1:621-1358       H--ANSSEPALLFRNIKCNYVFNNTLSRQLQP--------INYFDSYLGCVVNADNSTSS	123
BBM61037.1:621-1358       H--ANSSEPALLFRNIKCNYVFNNTLSRQLQP--------INYFDSYLGCVVNADNSTSS	123
BBM61397.1:621-1358       H--ANSSEPALLFRNIKCNYVFNNTLSRQLQP--------INYFDSYLGCVVNADNSTSS	123
BBM61267.1:621-1358       H--ANSSEPALLFRNIKCNYVFNNTLSRQLQP--------INYFDSYLGCVVNADNSTSS	123
P25192.1:621-1358         H--ANSSEPALLFRNIKCNYVFNNTLSRQLQP--------INYFDSYLGCVVNADNSTSS	123
ABD18914.1:621-1358       H--ANSSEPALLFRNIKCNYVFNNILSRQLQP--------INYFDSYLGCVVNADNSTSS	123
BBM61367.1:621-1358       H--ANSSEPALLFRNIKCNYVFNNTLSRQLQP--------INYFDSYLGCVVNADNSTSS	123
BBM61357.1:621-1358       H--ANSSEPALLFRNIKCNYVFNNTLSRQLQP--------INYFDSYLGCVVNADNSTSS	123
ABI93999.2:621-1358       H--ANSSEPASLFRNIKCNYVFNNTLSRQLQP--------INYFDSYLGCVVNADNSTSS	123
AZS64222.1:621-1358       H--ANSSEPALLFRNIKCNYVFNNTLLRQLQP--------INYFDSYLGCVVNADNSTSS	123
AZS64221.1:621-1358       H--ANSSEPALLFRNIKCNYVFNNTLLRQLQP--------INYFDSYLGCVVNADNSTSS	123
ANJ04974.1:621-1358       H--ANSSEPALLFRNIKCNYVFNNTLLRQLQP--------INYFDSYLGCVVNADNSTSS	123
BBM61337.1:621-1358       H--ANSSEPALLFRNIKCNYVFNNTLSRQLQP--------INYFDSYLGCVVNADNSTSS	123
AZS64223.1:621-1358       H--ANSSEPALLFRNIKCNYVFNNTLSRQLQP--------INYFDSYLGCVVNADNSTSS	123
QOV05164.1:621-1358       H--ANSSEPALLFRNIKCNYVFNNTLLRQLQP--------INYFDSYLGCVVNADNSTSS	123
QOV05143.1:621-1358       H--ANSSEPALLFRNIKCNYVFNNTLLRQLQP--------INYFDSYLGCVVNADNSTSS	123
QOV05147.1:621-1358       H--ANSSEPALLFRNIKCNYVFNNTLLRQLQP--------INYFDSYLGCVVNADNSTSS	123
QOV05175.1:621-1358       H--ANSSEPALLFRNIKCNYVFNNTLLRQLQP--------INYFDSYLGCVVNADNSTSS	123
ABD18916.1:621-1358       H--ANSSEPALLFRNIKCNYVFNNILSRQLQP--------INYFDSYLGCVVNADNSTSS	123
BBM61427.1:621-1358       H--ANSSEPALLFRNIKCNYVFNNTLSRQLQP--------INYFDSYLGCVVNADNSTSS	123
AAX38496.1:621-1358       H--ANSSEPALLFRNIKCNYVFNNILSRQLQP--------INYFDSYLGCVVNADNSTSS	123
AAX38492.1:621-1358       H--ANSSEPALLFRNIKCNYVFNNILSRQLQP--------INYFDSYLGCVVNADNSTSS	123
AAX38489.1:621-1358       H--ANSSEPALLFRNIKCNYVFNNILSRQLQP--------INYFDSYLGCVVNADNSTSS	123
NP_150077.1:621-1358      H--ANSSEPALLFRNIKCNYVFNNTLSRQLQP--------INYFDSYLGCVVNADNSTSS	123
BBM61347.1:621-1358       H--ANSSEPALLFRNIKCNYVFNNTLSRQLQP--------INYFDSYLGCVVNADNSTSS	123
BBM61217.1:621-1358       H--ANSSEPALLFRNIKCNYVFNNTLSRQLQP--------INYFDSYLGCVVNADNSTSS	123
BBM61527.1:621-1358       H--ANSSEPALLFRNIKCNYVFNNTLSRQLQP--------INYFDSYLGCVVNADNSTSS	123
BBM60947.1:621-1358       H--ANSSEPALLFRNIKCNYVFNNTLSRQLQP--------INYFDSYLGCVVNADNSTSS	123
BBM60957.1:621-1358       H--ANSSEPALLFRNIKCNYVFNNTLSRQLQP--------INYFDSYLGCVVNADNSTSS	123
ACJ66990.1:621-1358       H--ANSSEPALLFRNIKCNYVFNNTLSRQLQP--------INYFDSYLGCVVNADNSTSS	123
ACJ66961.1:621-1358       H--ANSSEPALLFRNIKCNYVFNNTLSRQLQP--------INYFDSYLGCVVNADNSTSS	123
ACJ66946.1:621-1358       H--ANSSEPALLFRNIKCNYVFNNTLSRQLQP--------INYFDSYLGCVVNADNSTSS	123
ACT10983.1:621-1358       H--ANSSEPALLFRNIKCNYVFNNTLSRQLQP--------INYFDSYLGCVVNADNSTSS	123
AAX38491.1:621-1358       H--ANSSEPALLFRNIKCNYVFNNTLSRQLQP--------INYFDSYLGCVVNADNSTSS	123
ABP38313.1:616-1353       H--ANSSEPALLFRNIKCNYVFNNTLSRQLQP--------INYFDSYLGCVVNADNSTSS	123
BBM61487.1:621-1358       H--ANSSEPALLFRNIKCNYVFNNTLSRQLQP--------INYFDSYLGCVVNADNSTSS	123
AAX38494.1:621-1358       H--ANSSEPALLFRNIKCNYVFNNTLSRQLQP--------INYFDSYLGCVVNADNSTSS	123
QBG67078.1:621-1358       H--ANSSEPALLFRNIKCNYVFNNTLSRQLQP--------INYFDSYLGCVVNADNSTSS	123
ABP38236.1:621-1358       H--ANSSEPALLFRNIKCNYVFNNTLSRQLQP--------INYFDSYLGCVVNADNSTSS	123
AVZ61119.1:621-1358       H--ANSSEPALLFRNIKCNYVFNNTLSRQLQP--------INYFDSYLGCVVNADNSTSS	123
Q9QAQ8.1:621-1358         H--ANSSEPALLFRNIKCNYVFNNTLSRQLQP--------INYFDSYLGCVVNADNSTSS	123
ABP38306.1:621-1358       H--ANSSEPALLFRNIKCNYVFNNTLSRQLQP--------INYFDSYLGCVVNADNSTSS	123
BBM61577.1:621-1358       H--ANSSEPALLFRNIKCNYVFNNTLSRQLQP--------INYFDSYLGCVVNADNSTSS	123
QBG67079.1:621-1358       H--ANSSEPALLFRNIKCNYVFNNTLSRQLQP--------INYFDSYLGCVVNADNSTSS	123
BBM60937.1:621-1358       H--ANSSEPALLFRNIKCNYVFNNTLSRQLQP--------INYFDSYLGCVVNADNSTSS	123
BBM61097.1:621-1358       H--ANSSEPALLFRNIKCNYVFNNTLSRQLQP--------INYFDSYLGCVVNADNSTSS	123
BBM61517.1:621-1358       H--ANSSEPALLFRNIKCNYVFNNTLSRQLQP--------INYFDSYLGCVVNADNSTSS	123
AVZ61109.1:621-1358       H--ANSSEPALLFRNIKCNYVFNNTLSRQLQP--------INYFDSYLGCVVNADNSTSS	123
BBM61077.1:621-1358       H--ANSSEPALLFRNIKCNYVFNNTLSRQLQP--------INYFDSYLGCVVNADNSTSS	123
BBM61317.1:621-1358       H--ANSSEPALLFRNIKCNYVFNNTLSRQLQP--------INYFDSYLGCVVNADNSTSS	123
BBM61567.1:621-1358       H--ANSSEPALLFRNIKCNYVFNNTLSRQLQP--------INYFDSYLGCVVNADNSTSS	123
BBM61457.1:621-1358       H--ANSSEPALLFRNIKCNYVFNNTLSRQLQP--------INYFDSYLGCVVNADNSTSS	123
                              .*.: * *:::::*. *       ** *         * * :  **:: *:: ..


SARS like

AGZ48806.1:3-1256       VITPGTNTSSEVAVLYQDVNCTDVPVAIHADQLTPSWRVYSTGNNVFQTQAGCLIGAEHV	641
ATO98132.1:3-1256       VITPGTNTSSEVAVLYQDVNCTDVPVAIHADQLTPSWRVYSTGNNVFQTQAGCLIGAEHV	641
ATO98218.1:3-1256       VITPGTNTSSEVAVLYQDVNCTDVPVAIHADQLTPAWRIYSTGNNVFQTQAGCLIGAEHV	641
ATO98231.1:3-1256       VITPGTNTSSEVAVLYQDVNCTDVPVAIHADQLTPAWRIYSTGNNVFQTQAGCLIGAEHV	641
AGZ48828.1:3-1256       VITPGTNTSSEVAVLYQDVNCTDVPVAIHADQLTPSWRVHSTGNNVFQTQAGCLIGAEHV	641
AGZ48818.1:3-1256       VITPGTNTSSEVAVLYQDVNCTDVPVAIHADQLTPSWRVYSTGNNVFQTQAGCLIGAEHV	641
ATO98157.1:1-1255       VITPGTNTSSEVAVLYQDVNCTDVPVAIHADQLTPAWRIYSTGNNVFQTQAGCLIGAEHV	642
ATO98205.1:1-1255       VITPGTNTSSEVAVLYQDVNCTDVPVAIHADQLTPSWRVYSTGNNVFQTQAGCLIGAEHV	642
AAV97990.1:1-1255       VITPGTNASSEVAVLYQDVNCTDVSTLIHADQLTPAWRIYSTGNNVFQTQAGCLIGAEHV	642
AAV98000.1:1-1255       VITPGTNASSEVAVLYQDVNCTDVSTLIHAEQLTPAWRIYSTGNNVFQTQAGCLIGAEHV	642
AAU04662.1:1-1255       VITPGTNASSEVAVLYQDVNCTDVSTLIHAEQLTPAWRIYSTGNNVFQTQAGCLIGAEHV	642
AAV91631.1:1-1255       VITPGTNASSEVAVLYQDVNCTDVSTLIHAEQLTPAWRIYSTGNNVFQTQAGCLIGAEHV	642
AAU04649.1:1-1255       VITPGTNASSEVAVLYQDVNCTDVSTLIHAEQLTPAWRIYSTGNNVFQTQAGCLIGAEHV	642
AAU04646.1:1-1255       VITPGTNASSEVAVLYQDVNCTDVSTLIHAEQLTPAWRIYSTGNNVFQTQAGCLIGAEHV	642
AAV98001.1:1-1255       VITPGTNASSEVAVLYQDVNCTDVSTLIHAEQLTPAWRIYSTGNNVFQTLAGCLIGAEHV	642
AAV97998.1:1-1255       VITPGTNASSEVAVLYQDVNCTDVSTLIHAEQLTPAWRIYSTGNNVFQTQAGCLIGAEHV	642
AAV49720.1:1-1255       VITPGTNASSEVAVLYQDVNCTDVSTLIHADQLTPAWRIYSTGNNVFQTQAGCLIGAEHV	642
AAV97985.1:1-1255       VITPGTNASSEVAVLYQDVNCTDVSTLIHAEQLTPAWRIYSTGNNVFQTQAGCLIGAEHV	642
AAU93319.1:1-1255       VITPGTNASSEVAVLYQDVNCTDVSTLIHAEQLTPAWRIYSTGNNVFQTQAGCLIGAEHV	642
AAV97992.1:1-1255       VITPGTNASSEVAVLYQDVNCTDVSTLIHAEQLTPAWRIYSTGNNVFQTQAGCLIGAEHV	642
AAV49722.1:1-1255       VITPGTNASSEVAVLYQDVNCTDVSTLIHAEQLTPAWRIYSTGNNVFQTQAGCLIGAEHV	642
AAU04664.1:1-1255       VITPGTNASSEVAVLYQDVNCTDVSTLIHAEQLTPAWRIYSTGNNVFQTQAGCLIGAEHV	642
AAV97989.1:1-1255       VITPGTNASSEVAVLYQDVNCTDVSTLIHAEQLTPAWRIYSTGNNVFQTQAGCLIGAEHV	642
AAS10463.1:1-1255       VITPGTNASSEVAVLYQDVNCTDVSTLIHAEQLTPAWRIYSTGNNVFQTQAGCLIGAEHV	642
AAV49723.1:1-1255       VITPGTNASSEVAVLYQDVNCTDVSTLIHAEQLTPAWRIYSTGNNVFQTQAGCLIGAEHV	642
AAV49719.1:1-1255       VITPGTNASSEVAVLYQDVNCTDVSTLIHAEQLTPAWRIYSTGNNVFQTQAGCLIGAEHV	642
AAV97986.1:1-1255       VITPGTNASSEVAVLYQDVNCTDVSTVIHAEQLTPAWRIYSTGNNVFQTQAGCLIGAEHV	642
AAV97984.1:1-1255       VITPGTNASSEVAVLYQDVNCTDVSTLIHAEQLTPAWRIYSTGNNVFQTQAGCLIGAEHV	642
AAV98002.1:1-1255       VITPGTNASSEVAVLYQDVNCTDVSTLIHAEQLTPAWRIYSTGNNVFQTQAGCLIGAEHV	642
AAU04661.1:1-1255       VITPGTNASSEVAVLYQDVNCTDVSTLIHAEQLTPAWRIYSTGNNVFQTQAGCLIGAEHV	642
AAV97995.1:1-1255       VITPGTNASSEVAVLYQDVNCTDVSTLIHAEQLTPAWRIYSTGNNVFQTQAGCLIGAEHV	642
AAS00003.1:1-1255       VITPGTNASSEVAVLYQDVNCTDVSTAIHADQLTPAWRIYSTGNNVFQTQAGCLIGAEHV	642
AAP51227.1:1-1255       VITPGTNASSEVAVLYQDVNCTDVSTAIHADQLTPAWRIYSTGNNVFQTQAGCLIGAEHV	642
AAP82968.1:1-1255       VITPGTNASSEVAVLYQDVNCTDVSTAIHADQLTPAWRIYSTGNNVFQTQAGCLIGAEHV	642
ABF68955.1:1-1255       VITPGTNASSEVAVLYQDVNCTDVSTAIHADQLTPAWRIYSTGNNVFQTQAGCLIGAEHV	642
AAR07631.1:1-1255       VITPGTNASSEVAVLYQDVNCTDVSTAIHADQLTPAWRIYSTGNNVFQTQAGCLIGAEHV	642
AAR07628.1:1-1255       VITPGTNASSEVAVLYQDVNCTDVSTAIHADQLTPAWRIYSTGNNVFQTQAGCLIGAEHV	642
AAR07627.1:1-1255       VITPGTNASSEVAVLYQDVNCTDVSTAIHADQLTPAWRIYSTGNNVFQTQAGCLIGAEHV	642
AAR07626.1:1-1255       VITPGTNASSEVAVLYQDVNCTDVSTAIHADQLTPAWRIYSTGNNVFQTQAGCLIGAEHV	642
AAR07629.1:1-1255       VITPGTNASSEVAVLYQDVNCTDVSTAIHADQLTPAWRIYSTGNNVFQTQAGCLIGAEHV	642
AAR07625.1:1-1255       VITPGTNASSEVAVLYQDVNCTDVSTAIHADQLTPAWRIYSTGNNVFQTQAGCLIGAEHV	642
ABF68958.1:1-1255       VITPGTNASSEVAVLYQDVNCTDVSTAIHADQLTPAWRIYSTGNNVFQTQAGCLIGAEHV	642
AAT74874.1:1-1255       VITPGTNASSEVAVLYQDVNCTDVSTAIHADQLTPAWRIYSTGNNVFQTQAGCLIGAEHV	642
AAR07624.1:1-1255       VITPGTNASSEVAVLYQDVNCTDVSTAIHADQLTPAWRIYSTGNNVFQTQAGCLIGAEHV	642
AAT76147.1:1-1255       VITPGTNASSEVAVLYQDVNCTDVSTAIHADQLTPAWRIYSTGNNVFQTQAGCLIGAEHV	642
AAR91586.1:1-1255       VITPGTNASSEVAVLYQDVNCTDVSTAIHADQLTPAWRIYSTGNNVFQTQAGCLIGAEHV	642
AAR07630.1:1-1255       VITPGTNASSEVAVLYQDVNCTDVSTAIHADQLTPAWRIYSTGNNVFQTQAGCLIGAEHV	642
AAP13567.1:1-1255       VITPGTNASSEVAVLYQDVNCTDVSTAIHADQLTPAWRIYSTGNNVFQTQAGCLIGAEHV	642
AAX16192.1:1-1255       VITPGTNASSEVAVLYQDVNCTDVSTAIHADQLTPAWRIYSTGNNVFQTQAGCLIGAEHV	642
ACZ71797.1:1-1255       VITPGTNASSEVAVLYQDVNCTDVSTAIHADQLTPAWRIYSTGNNAFQTQAGCLIGAEHV	642
ACZ71961.1:1-1255       VITPGTNASSEVAVLYQDVNCTDVSTAIHADQLTPAWRIYSTGNNVFQTQAGCLIGAEHV	642
ACZ71976.1:1-1255       VITPGTNASSEVAVLYQDVNCTDVSTAIHADQLTPAWRIYSTGNNVFQTQAGCLIGAEHV	642
ACZ72195.1:1-1255       VITPGTNASSEVAVLYQDVNCTDVSTAIHADQLTPAWRIYSTGNNVFQTQAGCLIGAEHV	642
ACZ72254.1:1-1255       VITPGTNASSEVAVLYQDVNCTDVSTAIHADQLTPAWRIYSTGNNVFQTQAGCLIGAEHV	642
ACZ72020.1:1-1255       VITPGTNASSEVAVLYQDVNCTDVSTAIHADQLTPAWRIYSTGNNVFQTQAGCLIGAEHV	642
ACZ71826.1:1-1255       VITPGTNASSEVAVLYQDVNCTDVSTAIHADQLTPAWRIYSTGNNVFQTQAGCLIGAEHV	642
ACZ72108.1:1-1255       VITPGTNASSEVAVLYQDVNCTDVSTAIHADQLTPAWRIYSTGNNVFQTQAGCLIGAEHV	642
ACB69905.1:1-1255       VITPGTNASSEVAVLYQDVNCTDVSTAIHADQLTPAWRIYSTGNNVFQTQAGCLIGAEHV	642
ACB69894.1:1-1255       VITPGTNASSEVAVLYQDVNCTDVSTAIHADQLTPAWRIYSTGNNVFQTQAGCLIGAEHV	642
ACB69883.1:1-1255       VITPGTNASSEVAVLYQDVNCTDVSTAIHADQLTPAWRIYSTGNNVFQTQAGCLIGAEHV	642
BAE93401.1:1-1255       VITPGTNASSEFAVLYQDVNCTDVSTAIHADQLTPAWRIYSTGNNVFQTQAGCLIGAEYV	642
AGT21078.1:1-1255       VITPGTNASSEVAVLYQDVNCTDVSTAIHADQLTPAWRIYSTGNNVFQTQAGCLIGAEYV	642
AFM43867.1:1-1255       VITPGTNASSEVAVLYQDVNCTDVSTAIHADQLTPAWRIYSTGNNVFQTQAGCLIGAEYV	642
ABF68957.1:1-1255       VITPGTNASSEVAVLYQDVNCTDVSTAIHADQLTPAWRIYSTGNNVFQTQAGCLIGAEHV	642
ABF68956.1:1-1255       VITPGTNASSEVAVLYQDVNCTDVSTAIHADQLTPAWRIYSTGNNVFQTQAGCLIGAEHV	642
ABF68959.1:1-1255       VITPGTNASSEVAVLYQDVNCTDVSTAIHADQLTPAWRIYSTGNNVFQTQAGCLIGAEHV	642
BAF42873.1:1-1255       VITPGTNASSEVAVLYQDVNCTDVSTAIHADQLTPAWRIYSTGNNVFQTQAGCLIGAEYV	642
AAR86775.1:1-1255       VITPGTNASSEVAVLYQDVNCTNVSAAIHADQLTPAWRIYSTGNNVFQTQAGCLIGAEHV	642
AEA10443.1:1-1255       VITPGTNASSEVAVLYQDVNCTDVSTAIHADQLTPAWRIYSTGNNVFQTQAGCLIGAEHV	642
ACQ82725.1:1-1255       VITPGTNASSEVAVLYQGVNCTDVSTAIHADQLTPAWRIYSTGNNVFQTQAGCLIGAEHV	642
AFR58714.1:1-1255       VITPGTNASSEVAVLYQDVNCTDVSTAIHADQLTPAWRIYSTGNNVFQTQAGCLIGAEHV	642
ACZ71991.1:1-1255       VITPGTNASSEVAVLYQDVNCTDVSTAIHADQLTPAWRIYSTGNNVFQTQAGCLIGAEHV	642
AEA10473.1:1-1255       VITPGTNASSEVAVLYQDVNCTDVSTAIHADQLTPAWRIYSTGNNVFQTQAGCLIGAEHV	642
ACZ72122.1:1-1255       VITPGTNASSEVAVLYQDVNCTDVSTAIHADQLTPAWRIYSTGNNVFQTQAGCLIGAEHV	642
ABD72995.1:1-1255       VITPGTNASSEVAVLYQDVNCTDVSTAIHADQLTPAWRIYSTGNNVFQTQAGCLIGAEHV	642
ABD72988.1:1-1255       VITPGTNASSEVAVLYQDVNCTDVSTAIHADQLTPAWRIYSTGNNVFQTQAGCLIGAEHV	642
ADC35483.1:1-1255       VITPGTNASSEVAVLYQDVNCTDVSTAIHADQLTPAWRIYSTGNNVFQTQAGCLIGAEHV	642
ABF65836.1:1-1255       VITPGTNASSEVAVLYQDVNCTDVSTAIHADQLTPAWRIYSTGNNVFQTQAGCLIGAEHV	642
AFR58672.1:1-1255       VITPGTNASSEVAVLYQDVNCTDVSTAIHADQLTPAWRIYSTGNNVFQTQAGCLIGAEHV	642
ACZ72093.1:1-1255       VITPGTNASSEVAVLYQDVNCTDVSTAIHADQLTPAWRIYSTGNNVFQTQAGCLIGAEHV	642
AAP33697.1:1-1255       VITPGTNASSEVAVLYQDVNCTDVSTAIHADQLTPAWRIYSTGNNVFQTQAGCLIGAEHV	642
ABD72969.1:1-1255       VITPGTNASSEVAVLYQDVNCTDVSTAIHADQLTPAWRIYSTGNNVFQTQAGCLIGAEHV	642
AAP41037.1:1-1255       VITPGTNASSEVAVLYQDVNCTDVSTAIHADQLTPAWRIYSTGNNVFQTQAGCLIGAEHV	642
ABD72977.1:1-1255       VITPGTNASSEVAVLYQDVNCTDVSTAIHADQLTPAWRIYSTGNNVFQTQAGCLIGAEHV	642
ABD72970.1:1-1255       VITPGTNASSEVAVLYQDVNCTDVSTAIHADQLTPAWRIYSTGNNVFQTQAGCLIGAEHV	642
AAU81608.1:1-1255       VITPGTNASSEVAVLYQDVNCTDVSTAIHADQLTPAWRIYSTGNNVFQTQAGCLIGAEHV	642
ABD73001.1:1-1255       VITPGTNASSEVAVLYQDVNCTDVSTAIHADQLTPAWRIYSTGNNVFQTQAGCLIGAEHV	642
AAR23250.1:1-1255       VITPGTNASSEVAVLYQDVNCTDVSTAIHADQLTPAWRIYSTGNNVFQTQAGCLIGAEHV	642
ABD72972.1:1-1255       VITPGTNASSEVAVLYQDVNCTDVSTAIHADQLTPAWRIYSTGNNVFQTQAGCLIGAEHV	642
ABD72985.1:1-1255       VITPGTNASSEVAVLYQDVNCTDVSTAIHADQLTPAWRIYSTGNNVFQTQAGCLIGAEHV	642
ABD72979.1:1-1255       VITPGTNASSEVAVLYQDVNCTDVSTAIHADQLTPAWRIYSTGNNVFQTQAGCLIGAEHV	642
ABD72984.1:1-1255       VITPGTNASSEVAVLYQDVNCTDVSTAIHADQLTPAWRIYSTGNNVFQTQAGCLIGAEHV	642
P59594.1:1-1255         VITPGTNASSEVAVLYQDVNCTDVSTAIHADQLTPAWRIYSTGNNVFQTQAGCLIGAEHV	642
ABD72982.1:28-1255      VITPGTNASSEVAVLYQDVNCTDVSTAIHADQLTPAWRIYSTGNNVFQTQAGCLIGAEHV	615
QND76020.1:5-1259       VITPGTNASSEVAVLYQDVNCTDVPTAIRADQLTPAWRVYSTGVNVFQTQAGCLIGAEHV	642
QND76034.1:5-1259       VITPGTNASSEVAVLYQDVNCTDVPTAIRADQLTPAWRVYSTGVNVFQTQAGCLIGAEHV	642
AVP78042.1:12-1245      VITPGTNTSSEVAVLYQDVNCTDVPTTIHADQLTPAWRIYAIGTSVFQTQAGCLIGAEHV	621
AVP78031.1:12-1246      VITPGTNTSLEVAVLYQDVNCTDVPTTIHADQLTPAWRIYATGTNVFQTQAGCLIGAEHV	622
REF                     VITPGTNTSNQVAVLYQDVNCTEVPVAIHADQLTPTWRVYSTGSNVFQTRAGCLIGAEHV	656
QHR63300.2:1-1269       VITPGTNASNQVAVLYQDVNCTEVPVAIHADQLTPTWRVYSTGSNVFQTRAGCLIGAEHV	656
6ZGF_A:32-1252          VITPGTNTSNQVAVLYQDVNCTEVPVAIHADQLTPTWRVYSTGSNVFQTRAGCLIGAEHV	656
                        *******:* :.*****.****:* . *:*:****:**::: * ..*** ********:*
```
